# GGA1 interacts with the endosomal Na+/H+ Exchanger NHE6 governing localization to the endosome compartment

**DOI:** 10.1101/2023.11.08.565997

**Authors:** Li Ma, Ravi Kiran Kasula, Qing Ouyang, Michael Schmidt, Eric M. Morrow

## Abstract

Mutations in the endosomal Na+/H+ exchanger (NHE6) cause Christianson syndrome (CS), an X-linked neurological disorder. Previous studies have shown that NHE6 functions in regulation of endosome acidification and maturation in neurons. Using yeast two-hybrid screening with the NHE6 carboxyl-terminus as bait, we identify Golgi-associated, Gamma adaptin ear containing, ARF binding protein 1 (GGA1) as an interacting partner for NHE6. We corroborated the NHE6-GGA1 interaction using co-immunoprecipitation (co-IP): using over-expressed constructs in mammalian cells; and co-IP of endogenously-expressed GGA1 and NHE6 from neuroblastoma cells, as well as from mouse brain. We demonstrate that GGA1 interacts with organellar NHEs (NHE6, NHE7 and NHE9) but not with cell-surface localized NHEs (NHE1 and NHE5). By constructing hybrid NHE1/NHE6 exchangers, we demonstrate that the cytoplasmic tail of NHE6 is necessary and sufficient for interactions with GGA1. We demonstrate the co-localization of NHE6 and GGA1 in cultured, primary hippocampal neurons, using super-resolution microscopy. We test the hypothesis that the interaction of NHE6 and GGA1 functions in the localization of NHE6 to the endosome compartment. Using subcellular fractionation experiments, we show that NHE6 is mis-localized in GGA1 knockout cells wherein we find less NHE6 in endosomes but more NHE6 transport to lysosomes, and more Golgi retention of NHE6 with increased exocytosis to the surface plasma membrane. Consistent with NHE6 mis-localization, and Golgi retention, we find the intra-luminal pH in Golgi to be alkalinized. Our study demonstrates a new interaction between NHE6 and GGA1 which functions in the localization of this intra-cellular NHE to the endosome compartment.

## INTRODUCTION

Loss-of-function mutations in the X-linked endosomal Na+/H+ Exchanger 6 (NHE6, encoded by the SLC9A6 gene) cause the neurological disorder Christianson Syndrome (CS) (1–10). In mammalian cells, NHE6, NHE7, NHE8 and NHE9 are generally considered to be intra-cellular, with NHE6 and NHE9 studied as endosomal NHEs (4,11–15). On the other hand, NHE1, NHE2, NHE3, NHE4 and NHE5 are reported to be cell-surface NHEs (15,16). NHEs are evolutionary-conserved transmembrane proteins that regulate the electroneutral exchange of H+ ions with Na+ or K+ ions (17,18). NHE proteins are composed of a conserved 12 membrane-spanning domain in the N-terminus with a Na+/H+ exchanger, and a less-conserved cytoplasmic C-terminus that is involved in protein binding (14). They regulate a range of critical physiological functions including cytosolic/organellar pH, cell volume, acid-base homeostasis as well as broader cellular functions, such as intracellular trafficking and posttranslational modification (14). Mutations of some plasma membrane NHEs are associated with neural dysfunction (14,19–25). Also, NHE5 has been shown to modulate synaptic plasticity by negatively regulating activity-dependent dendritic spine growth (26). Prior studies have demonstrated that loss of NHE6 leads to over-acidification of endosomes, as well as defects in endosome maturation and lysosome function (4,8,27); however, the cellular mechanisms of disease in CS, and the function of NHE6 in endosome maturation are incompletely understood.

Protein-protein interactions provide important clues for cellular functions; however, very few NHE6-interacting proteins are known. Cytosolic domains of the endosomal NHE6 and NHE9 are reported to bind the scaffold protein RACK1, and the NHE6-Rack1 interaction is proposed to control receptor recycling in cultured cells (28). NHE6 also directly interacts with secretory carrier membrane protein 5 (SCAMP5), and this SCAMP5-dependent recruitment of NHE6 to synaptic vesicles plays a critical role in manifesting presynaptic efficacy both at rest and during synaptic plasticity (29,30).

Here, we used yeast two-hybrid screening to find new NHE6 interacting partners. We found that GGA1, a member of the Golgi-localized γ-ear-containing ADP-ribosylation factor (ARF)-binding (GGA) protein family (31–41), is a new NHE6-interacting protein. The NHE6 cytoplasmic domain plays the main role in GGA1 interaction. GGA1 also binds strongly to other intracellularly-localized NHEs, including NHE7 and NHE9, but less or not at all to plasma membrane localized NHE1 and NHE5. GGA proteins are monomeric clathrin adaptors that package and traffic cargo from the trans-Golgi network (TGN) to the endocytic pathway, and also mediate retrograde trafficking from the endocytic pathway to the TGN (34,36,42–48). There are three members of the mammalian GGA family (GGA1-3), that all contain: (1) an N-terminal VHS domain that binds proteins with an acidic di-leucine signal like mannose 6-phosphate receptors (M6PRs), (2) a GAT domain that binds Arf:GTP, (3) a connecting hinge region that recruits clathrin, and (4) a C-terminal GAE domain that recruits accessory proteins (45,49). GGAs traffic cation-dependent and -independent M6PRs, which are critical for delivering newly-synthesized precursor lysosomal enzymes to the endocytic pathway where they will ultimately participate in the degradation of cellular materials in acidic lysosomes (38,41,48,50–52). Here we demonstrate that the interaction of GGA1 with NHE6 functions in the localization of NHE6 to the endosome compartment, and in the absence of GGA1, NHE6 is mislocalized including retention in the Golgi leading to intra-lumen alkalinization and increased exocytosis of NHE6 to the cell surface membrane.

## RESULTS

### Identification of GGA1 as an NHE6-interacting partner

A yeast two-hybrid screen was performed using the mouse NHE6 cytoplasmic tail amino acids Leu^537^-Asp^630^ (NP_766368.2) as bait to discover possible NHE6-interacting proteins from a random-primed rat hippocampus cDNA library. A total of 53.8 million cDNA clones were screened and 123 His+ colonies were selected on DO-3 medium lacking tryptophan, leucine, and histidine. As shown in Figure 1A, NHE6 with either GGA1 (lane 5) or Septin8 (Sept8, lane 8) in bait-prey combination allowed growth on DO-3 medium, while negative controls containing empty bait and prey vectors, pB27 and pP7 (lane 2), or pB27 and pP7 in combination with prey and bait molecules (lane 3, 4, 6 and 7) did not grow on this medium. Among these 123 colonies, sequence analyses revealed that 13 coded for GGA1 (Table S1). All GGA1 sequences showed the maximally high confidence Predicted Biological Score (PBS). Another 24 clones coded for Sept8 which also showed very high confidence scores; however, Sept8 failed to show an interaction with NHE6 in the following immunoprecipitation (IP) analysis and was not studied further. These results from yeast two-hybrid screening suggest that GGA1 is a new interaction partner for NHE6.

**Figure 1.**
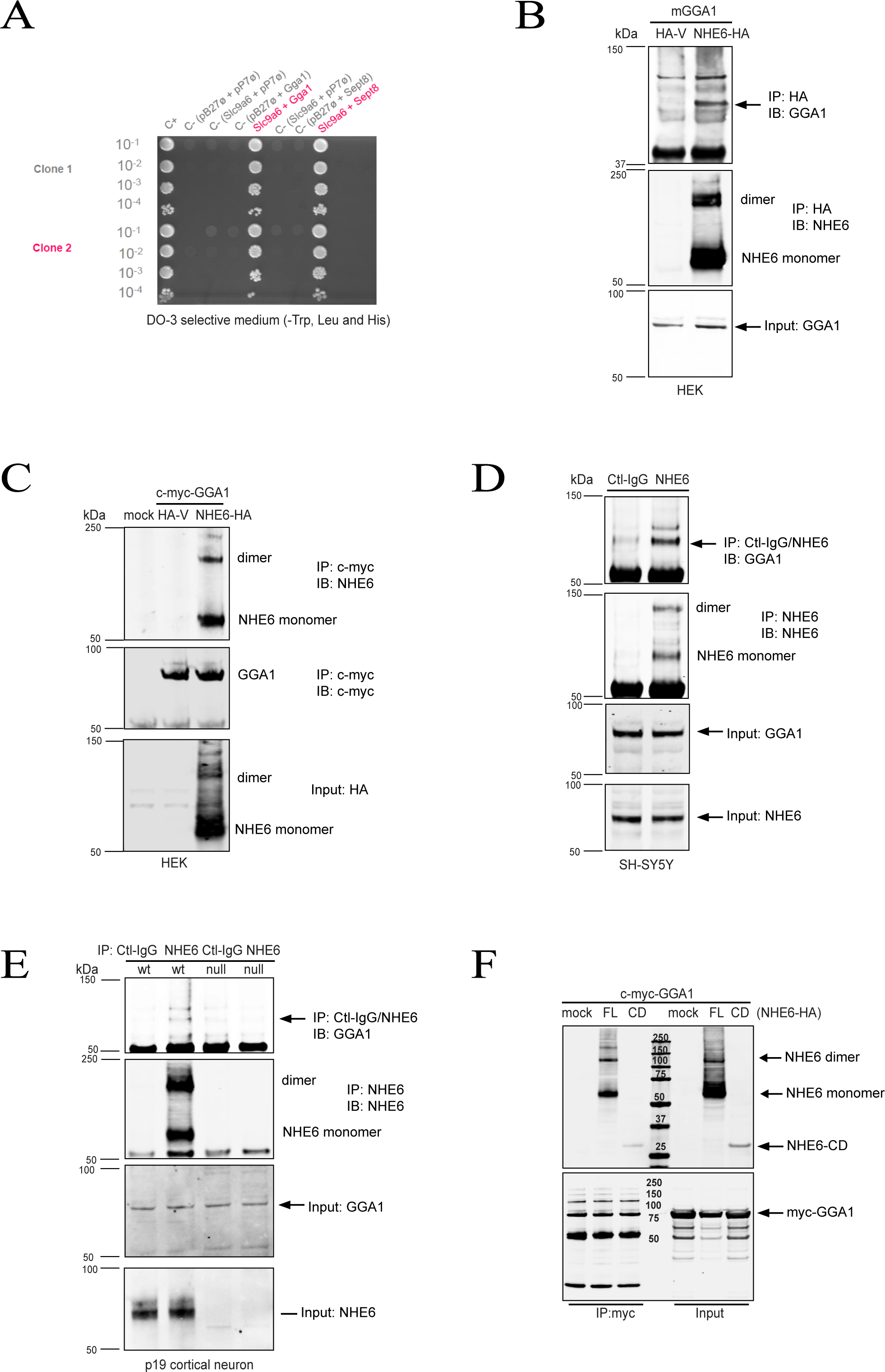
Identification of GGA1 as a new NHE6 interacting partner. **(A)** Yeast cells containing bait construct pB27-NHE6 (N-LexA-NHE6-537-630, amino acids 537-630 of *Slc9a6* cloned into the pB27 plasmid hgx2030v2_pB27) and prey constructs: pP6-GGA1 (N-GAL4-GGA1, clone RHC_RP_hgx2030v2_pB27_C-38 cloned into pP6) or pP6-Sept8 (N-GAL4-Sept8, clone RHC_RP_hgx2030v2_pB27_B-2 cloned into pP6). Yeast cells were obtained by mating and spotted, at the dilutions indicated, on DO-3 selective media lacking Trp, Leu and His. Negative controls included the following empty bait or pray vectors: pB27 and pP7, or pB27+Gga1, pB27+Sept8, Slc9a6+pP7. **(B)** Cell lysates from HEK293T cells expressing mGGA1 with either NHE6-HA or HA-Vector were immunoprecipitated with anti-HA antibody. The precipitates were analyzed with both anti-GGA1 and anti-NHE6 blotting. Total cell lysates (TCLs) were subjected to western blot analysis with anti-GGA1 antibody. All samples were separated by 4-12% SDS–PAGE unless mentioned separately. **(C)** Cell lysates from HEK293T cells expressing c-Myc-GGA1 with either NHE6-HA or HA-Vector were immunoprecipitated with anti-c-Myc antibody. The precipitates were probed with anti-NHE6 and c-Myc antibodies. TCLs were subjected to western blot analysis with anti-HA to detect the expression of NHE6. **(D)** Co-IP assay in SH-SY5Y neuroblastoma cells to detect interaction between endogenous NHE6 and GGA1. TCLs from SH-SY5Y cells were precipitated by anti-NHE6 antibody, with normal rabbit IgG used as a control. The precipitates were subjected to western blot analysis using anti-GGA1 and anti-NHE6 antibodies. TCLs were subjected to western blot analysis with anti-NHE6 and anti-GGA1 to detect the expression level of endogenous NHE6 and GGA1. **(E)** Brain tissue from NHE6-null and wildtype (WT) males at postnatal day 19 (P19) was collected and homogenized. TCLs were precipitated by anti-NHE6 antibody, with normal rabbit IgG used as a control. The precipitates were subjected to western blot analysis using both anti-GGA1 and anti-NHE6 antibodies. TCLs were subjected to western blot analysis with anti-NHE6 and anti-GGA1 antibodies to detect the expression level of GGA1 and NHE6. **(F)** TCLs from HEK293T cells expressing c-Myc-GGA1 with either HA-tagged full-length NHE6 (NHE6-FL) or cytoplasmic domain NHE6 (NHE6-CD) were immunoprecipitated with anti-c-Myc antibody. The precipitates were probed with HA and c-Myc antibodies. TCLs were subjected to western blot analysis with anti-HA or c-Myc antibodies to detect the expression of NHE6-FL or NHE6-CD and myc-GGA1.

IP analysis was then used to confirm this NHE6-GGA1 interaction in mammalian cells. To test whether NHE6 interacts with GGA1 in mammalian cells, HEK293T cells were co-transfected with pCMV-mGGA1 and either pCDNA3.1/NHE6-HA or HA-vector. Cell lysates were immunoprecipitated with anti-HA antibody to pull down NHE6 and subjected to western blot analysis to detect bound GGA1 using anti-GGA1 antibody. As shown in Figure 1B, GGA1 was detected in anti-HA immunoprecipitates from cells expressing GGA1 and NHE6, but not from cells expressing GGA1 and HA vector, indicating NHE6 is able to pull down and interact with GGA1.

We then tested whether GGA1 could pull down NHE6. HEK293T cells were co-transfected with pCDNA3.1/c-Myc-GGA1 and either pCDNA3.1/NHE6-HA or HA-vector. Cell lysates were immunoprecipitated with anti-c-Myc antibody to pull down GGA1 and subjected to western blot analysis to detect bound NHE6 using a custom-made anti-NHE6 antibody (4). As shown in Figure 1C, NHE6 was detected in anti-c-Myc immunoprecipitates from cells expressing GGA1 and NHE6, but not from cells expressing GGA1 and HA-vector, indicating GGA1 is able to co-precipitate and interact with NHE6.

To examine whether this interaction between NHE6 and GGA1 could be detected endogenously without over-expression of constructs, SH-SY5Y neuroblastoma cell lysates were precipitated with either anti-NHE6 antibody or control-IgG, followed by immunoblotting with anti-GGA1 antibody. GGA1 was detected in immunoprecipitates from NHE6 precipitates, but not control-IgG precipitates (Figure 1D). This result supports a physical interaction between NHE6 and GGA1 in endogenously expressed proteins.

Since NHE6 is highly expressed in the brain (4), we then investigated the NHE6-GGA1 interaction in cortical brain tissue from NHE6-null and wildtype (WT) littermate controls at postnatal day 19 (P19). Homogenized cortical brain lysates were immunoprecipitated with anti-NHE6 antibody or control-IgG, followed by immunoblotting with anti-GGA1 antibody. In WT cortical tissue, GGA1 was detected in anti-NHE6 precipitates, but not control-IgG precipitates (Figure 1E). Further, GGA1 was not detected in NHE6-null tissue (Figure 1E). There were no differences in GGA1 protein expression between NHE6-null and WT brain tissue(Figure 1E, 3^rd^ panel). This result further demonstrates that NHE6 interacts with GGA1in mouse brain tissue.

To further confirm the NHE6 carboxyl-terminus alone can interact with GGA1 in mammalian cells, HEK293T cells were co-transfected with pCDNA3.1/c-Myc-GGA1 and either a full-length NHE6 (pCDNA3.1/NHE6-FL-HA) or cytoplasmic domain (CD, pCDNA3.1/NHE6-CD-HA) construct. Cell lysates were immunoprecipitated with anti-Myc antibody to pull down GGA1 and then subjected to western blot analysis to detect bound NHE6 using anti-HA antibody. Both full-length and CD NHE6 products were detected in anti-Myc immunoprecipitated, indicating the NHE6 cytoplasmic domain alone can interact with GGA1 (Figure 1F).

### GGA1 shows strong interaction with organellar NHEs (NHE6/7/9) but not with cell surface localized NHEs (NHE1/5)

To determine whether GGA1 solely interacts with NHE6 or interacts with other NHEs, we tested the extent to which plasma membrane (e.g. NHE5) and organellar (NHE7 and NHE9) NHEs interact with GGA1 (Figure B and S1A). NHE5, NHE7 and NHE9 were all constructed with an HA-tag in the pCDNA3.1 vector, and expression was confirmed by both HA (Figure S1A) and NHE-specific antibodies (Figure S1B). We then immunoprecipitated HEK293T cells that were co-transfected with pCDNA3.1/c-Myc-GGA1 and HA-tagged NHEs (NHE5, NHE6, NHE7 or NHE9). Like NHE6 (Figure S1C, top panel, lane2), organellar NHE7 and NHE9 (Figure S1C, top panel, lanes 4 and 5) interact with GGA1. The plasma membrane NHE5 had dramatically reduced potential for interaction with GGA1 (Figure S1C, top panel, lane 3). This indicates that GGA1 preferentially interacts with organellar NHEs compared to the plasma membrane NHE5.

We then examined whether GGA1 interacts with the plasma membrane localized NHE1. We were unable to detect any interaction between GGA1 and NHE1 (Figure 2A). To further investigate the NHE domain of interaction with GGA1, we then generated two NHE6-NHE1 chimeric constructs by using the in-fusion snap method (Takara), that swaps the N-terminus exchanger domain and C-term cytoplasmic tail between NHE1 and NHE6 (Figure 2B). As shown in Figure 2C, the NHE6 C-terminus with the NHE1 N-terminus (NHE1N/NHE6C) maintains a strong interaction with GGA1. However, the NHE1 C-terminus with the NHE6 N-terminus (NHE6N/NHE1C) abolishes the interaction with GGA1 (Figure 2C). These findings support the interpretation that the NHE6 C-term is both sufficient and necessary for the interaction with GGA1.

**Figure 2.**
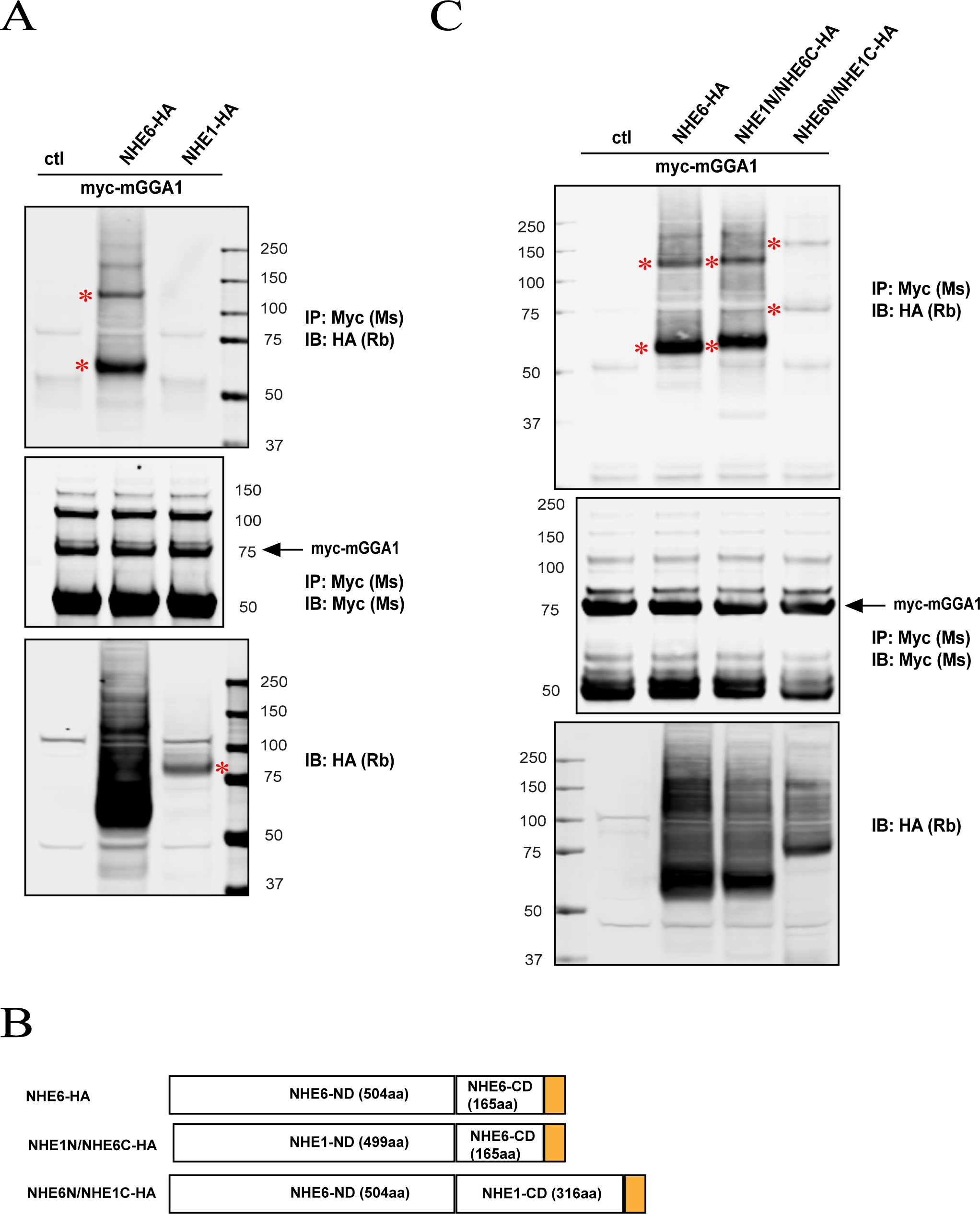
GGA1 interacts with both NHE6 N- and C-terminus domains and not NHE1. **(A)** Total cell lysates (TCLs) from HEK293T cells expressing c-Myc-GGA1 with either NHE6-HA or NHE1-HA were immunoprecipitated with anti-c-Myc antibody. The precipitates were probed with HA and c-Myc antibodies. TCLs were subjected to western blot analysis with the anti-HA antibody to detect the expression of NHE6 and NHE1. **(B)** Domain schematics of HA-tagged NHE6, and chimeric snap constructs NHE1-N terminus+NHE6-C terminus (NHE1N/NHE6C) and NHE6-N terminus+NHE1-C terminus (NHE6N/NHE1C). **(C)** TCLs from HEK293T cells expressing c-Myc-GGA1 with HA-tagged NHE6, NHE1N/NHE6C and NHE6N/NHE1C were immunoprecipitated with anti-c-Myc antibody. The precipitates were probed with HA and c-Myc antibodies. TCLs were subjected to western blot analysis with anti-HA to detect the expression of NHE6 and chimeric snap constructs.

To determine the extent to which NHE interaction is unique to GGA1, we performed IP experiments in GGA3. Specifically, we pulled down GGA3 and examined its interaction with either NHE6 or NHE9 in co-expression studies in HEK293T cells. Interestingly, as shown from Figure S2A, both NHE6 and NHE9 were co-immunoprecipitated by GGA3 (lane 2 and 3). Also, GGA3 and GGA1 could form a heterodimer (lane 4). Thus, our results suggest a broad interaction between GGAs and endosomal NHE family members.

### GGA1 GAE domain shows the strongest interaction with NHE6

GGA1 is composed of a VHS domain, GAT domain, GAE domain and linker Hinge domain in between GAT and GAE (31,32,53–55). To identify the region of GGA1 that mediates the interaction with NHE6, we constructed a series of domains (VHS, GAT, GAE and Hinge) (Figure 3A) to compare the ability to bind NHE6 by co-IP assay. As shown in Figure 2B, all GGA1 single domains, except the Hinge domain, interact with NHE6 (Figure 3C). The GAE domain showed the strongest binding activity with NHE6 (Figure 3C). To confirm that the Hinge domain does not interact with NHE6, we constructed more GGA1 domains (delGAE, delVHS, VHS+GAT, Hinge+GAE, GAT+Hinge and delHinge). All of these domains interacted with NHE6, and deletion of the Hinge region (delHinge) showed the strongest interaction with NHE6 (Figure 3D and Figure 3E). Therefore, the Hinge domain may function as dominant negative domain wherein this domain weakens the GGA1-NHE6 interaction. Compared to delGAE (VHS+GAT+Hinge), VHS+GAT has a stronger binding activity further supporting the Hinge linker’s dominant negative function. Although GAE shows the strong binding activity (Figure 3B and 3C), Hinge+GAE dramatically reduced the binding activity, further indicating that the Hinge domain functions as a dominant negative.

**Figure 3.**
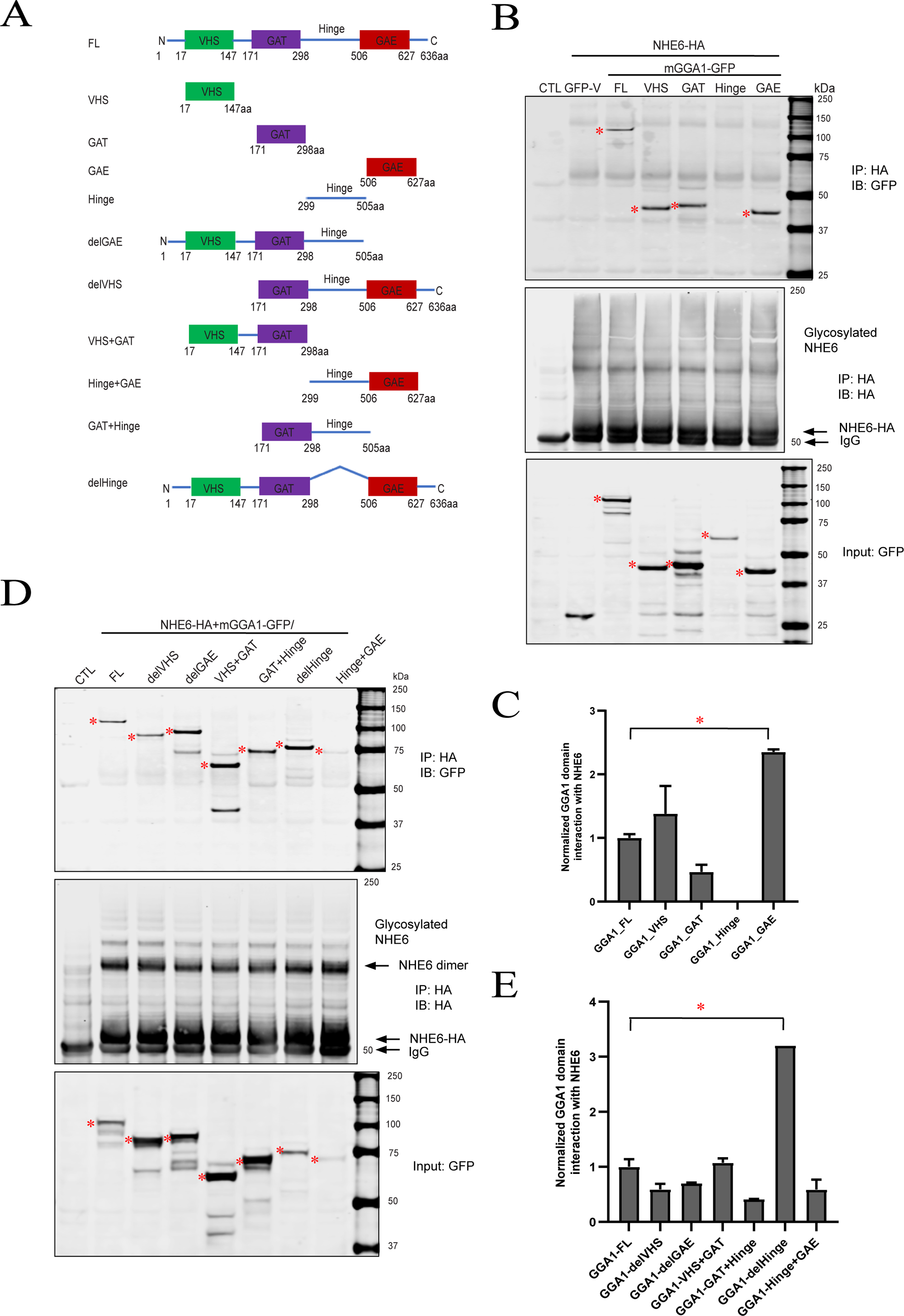
Mapping of interaction regions between NHE6 and GGA1. **(A)** Structure of mGGA1 domains: VHS (aa(17–147), green), GAT (aa(171–298), purple), GAE (aa(506–627), red), Hinge (aa(299–505)), del GAE (aa(1–505)), delVHS (aa(171–636)), VHS+GAT (aa(17–298)), Hinge+GAE (aa(299–627)), GAT+Hinge (aa(171–505)), and del Hinge (del aa(299–505)). **(B, D)** GFP-tagged GGA1 construct mGGA1-FL/VHS/GAT/Hinge/GAE (**B**), or GFP-tagged GGA1 construct mGGA1-FL/delVHS/delGAE/VHS+GAT/GAT+Hinge/delHinge/Hinge+GAE (**D**) was co-transfected with NHE6-HA in HEK293T cells. Anti-HA immunoprecipitates were analyzed with anti-GFP immunoblotting to identify GGA1 region(s) necessary for NHE6 interaction. TCLs were subjected to western blot analysis with anti-GFP to detect the expression level of GGA1 constructs. **(C, E)** GGA1 domain interaction with NHE6 was calculated by normalization of immunoprecipitated products with corresponding inputs. (mean±SEM; n=2; *p<0.05, unpaired *t* test)

### Co-localization of NHE6 and GGA1 in neurons using super-resolution microscopy

The finding that NHE6 interacts with GGA1 led us to explore where the co-localization of these two proteins occurs in cells such as neurons. NHE6 mainly localizes in early, recycling and late endosomes (4,12,56). In dissociated hippocampal neurons *in vitro*, NHE6 localization is punctate, adjacent to Golgi apparatus and distributed throughout the cell (4). Mouse hippocampal neurons were stained for NHE6 and GGA1 using immunocytochemistry and imaged using structured illumination microscopy (SIM) (Figure 4A). As shown in Figure 4A, NHE6 and GGA1 colocalize in the perinuclear region. Further Pearson’s colocalization analysis showed a positive correlation (Pearson correlation coefficient=0.35) (Figure 4B).

**Figure 4.**
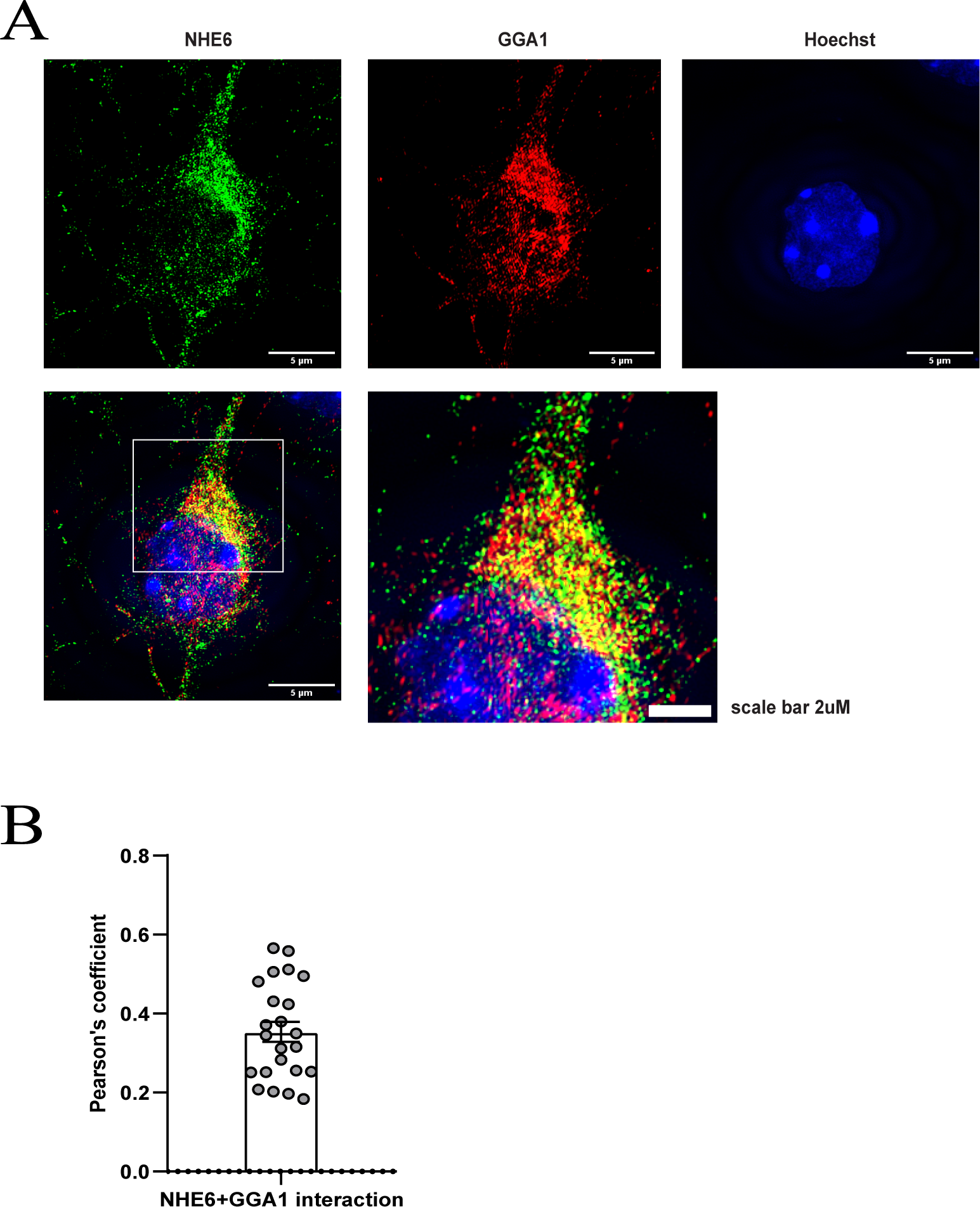
NHE6 and GGA1 colocalization using structured illumination microscopy (SIM) in mouse neurons *in vitro*. **(A)** Representative image of mouse primary hippocampal neurons at 3 days *in vitro* (3 DIV) were co-stained with anti-NHE6 (green) and anti-GGA1 (red) antibodies. Hoechst=blue. Scale bar=5 μm, Inset scale bar=2uM. **(B)** Quantification of NHE6-GGA1 colocalization by Pearson correlation coefficient analysis (n=23 cells from 3 unique neuronal cultures, mean±SEM).

We also used a 5x expansion microscopy technique (57) to acquire multicolor z-stack super-resolution images of rat hippocampal neurons, co-labelled with NHE6, GGA1, and the Golgi marker Giantin (Figure 5A, B). We created surfaces for Giantin and spots for NHE6 and GGA1 (Figure 5C, D) for quantification. From the shortest distance of these spots, and their proximity to Giantin surfaces, we classified the NHE6 and GGA1 spots into six different categories (Figure 5C, D). Expansion factor is measured, and the average shortest distance values are shown in Figure S3. Based on the number of spots in each category, we found that a fraction of 0.2084± 0.06840 (mean±SD) GGA1 spots co-localize with NHE6, of which 0.04189±0.03159 (mean±SD) NHE6/GGA1 co-labelled spots are within Giantin (Figure 5E). About 0.2412±0.08785 (mean±SD) NHE6 spots co-localize with GGA1 out of which 0.05129±0.04874 (mean±SD) NHE6/GGA1 co-labelled spots are within Giantin (Figure 5E). These expansion results further confirm NHE6 and GGA1 colocalization using microscopy techniques.

**Figure 5.**
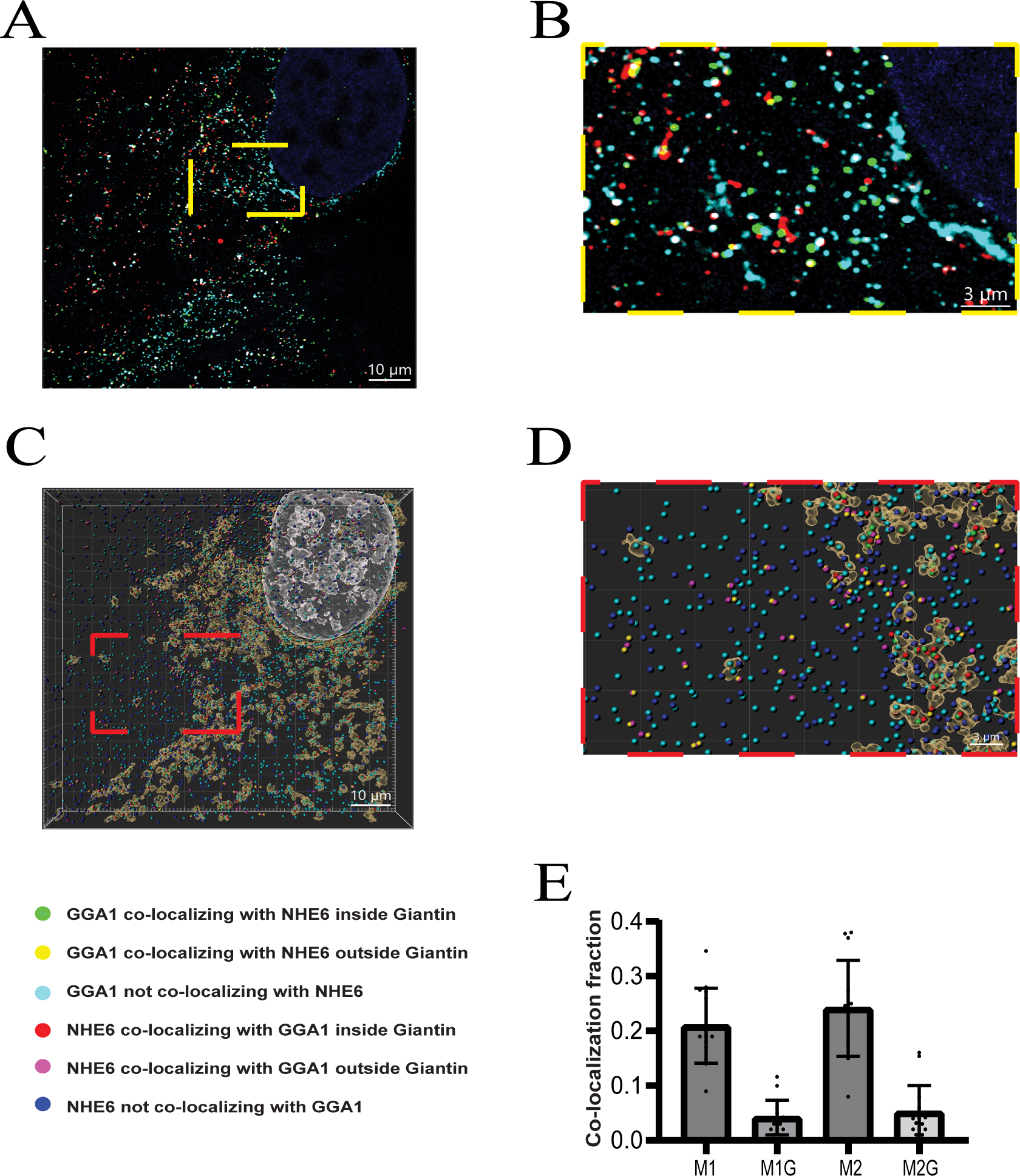
Colocalization of NHE6 and GGA1 in Golgi by 5x expansion microscopy imaging. Co-localization of GGA1 and NHE6 with respect to Golgi in rat hippocampal neurons: (**A**) shows a single frame of 5x expanded rat hippocampal neuron on DIV14 co-stained for nucleus (blue), Giantin (cyan), GGA1 (red), and NHE6 (green) from a z-stack acquired using confocal microscopy. Corrected scale bar: ∼2.27 uM. The intensities were adjusted for visualization. **(B)** enlarged image from **(A)** shown in yellow box. **(C)** Shows the 3D re-construction of the z-stack using IMARIS surface tools for the nucleus (white), and Giantin (brown) and spot classification for GGA1 (green, yellow, or cyan), and NHE6 (red, pink, or blue). The spots are classified based on the co-localization of a GGA1 or NHE6 with the other stain. Corrected scale bar: ∼2.27 uM. **(D)** enlarged image from **(C)** shown in red box. **(E)** Co-localization fraction of GGA1 or NHE6 analyzed using IMARIS software. M1: fraction of GGA1 co-localizing with NHE6; M1G: fraction of co-localizing GGA1 with NHE6 inside Giantin; M2: fraction of NHE6 co-localizing with GGA1; M2G: fraction of co-localizing NHE6 with GGA1 inside Giantin. Corrected scale bar: ∼0.68 uM. The color code for **(C)** and **(D)** are described to the right of image. (n=14 cells from 4 unique neuronal cultures).

### GGA1 functions to stabilize NHE6 in the endosome compartment

To elucidate the functional importance of the interaction between NHE6 and GGA1, we established two GGA1-null subclonal cell lines that were gene-edited to knockout GGA1 (Table S2). Sanger sequencing confirm the gene targeting events in the two lines (Figure S4A-C). Further western blotting (Figure S5A) shows these GGA1-null lines lead to the loss of GGA1 protein. Protein expression of GGA3 and tubulin was not changed (Figure S5A), demonstrating the specificity of our GGA1 knockout lines. Also, NHE6 protein level was not altered, suggesting that loss of GGA1 does not affect NHE6 protein levels (Figure S5A).

Accumulating evidence indicates that GGAs localize to Golgi and endosomes and regulate cargo trafficking between the Golgi and endosomes (33,34,53,58–60). NHE6 is distributed throughout the endocytic pathway, (4,12,61). Thus, we hypothesize that GGA1 regulates NHE6 trafficking from both Golgi to endosomes and retrograde endosome-Golgi. To test this hypothesis, we conducted a series of subcellular fractionations to investigate NHE6 distribution across different organelles in the presence and absence of GGA1 using HAP1-GGA1-KO cell lines. HAP1-WT and GGA1-KO cells were used to fractionate endosomes using a method based on protocols published by de Araujo et al (2015a and b) (62,63). The purity of the fractionation product was verified by detecting different organelle markers: Rab5 for early endosomes (EE), Rab7 for late endosomes (LE), Rab11 for recycling endosomes (RE); LAMP1 for lysosomes; and GM130 for Golgi (Figure 6A). As shown from Figure 6B, GGA1 KO lines had a significant reduction of NHE6 in the endosome fraction, indicating GGA1 is playing a role to stabilize NHE6 in endosomes. We then investigated how loss of GGA1 affected NHE6 using a fractionation method to isolate a cellular fractionation enriched for the late endosome compartment (62,63). We observed a 15% reduction of NHE6 in the late endosome compartment in GGA1 KO lines compared to control cells (Figure 6C, D). These data support the interpretation that GGA1 is required to stabilize NHE6 localization to endosome compartments.

**Figure 6.**
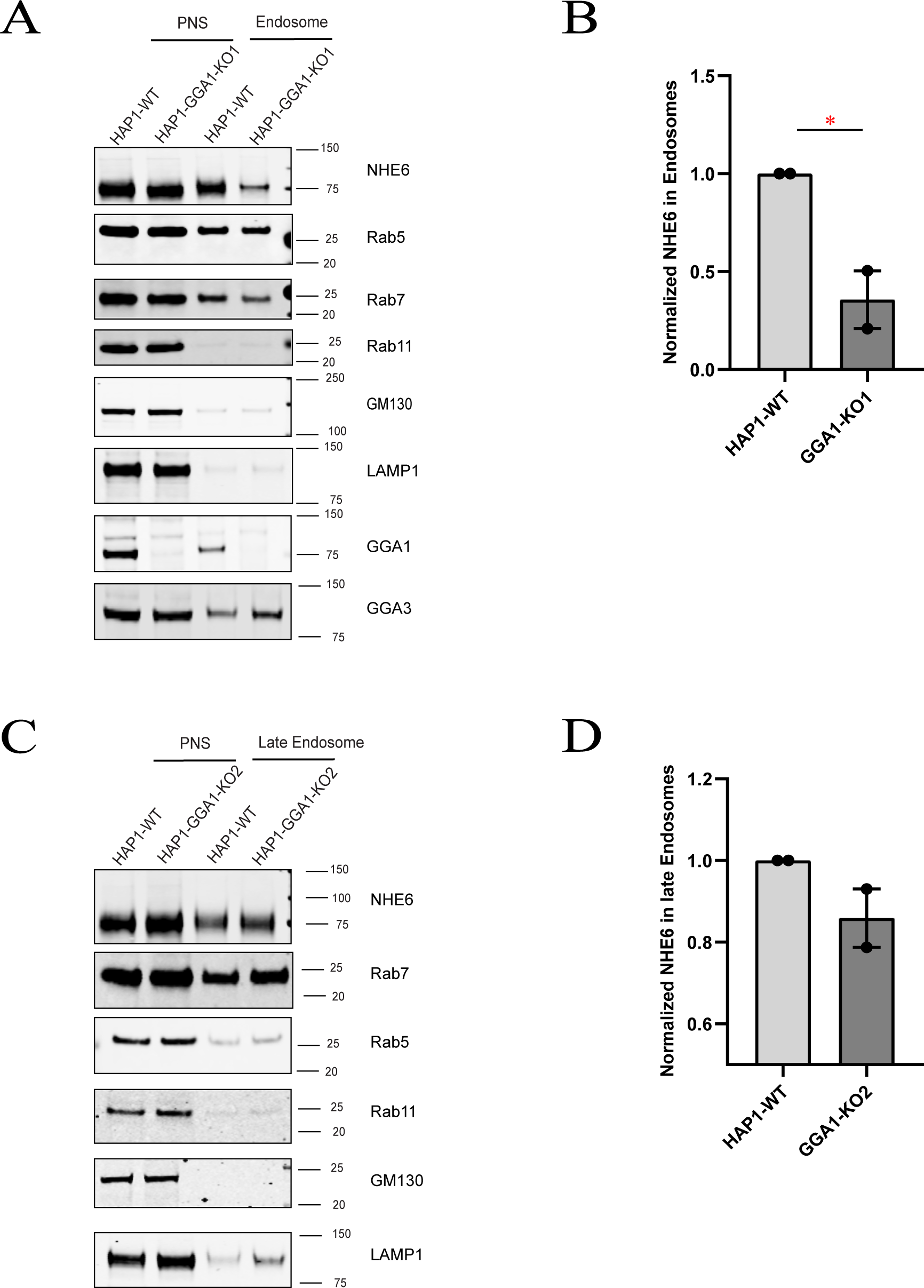
Loss of GGA1 leads to less NHE6 in endosomes. **(A)** Western blot of NHE6 protein in early endosome fractionation in HAP1 GGA1 knockout (KO) line 1 and wildtype (WT) cells using the following markers: Rab5 (early endosome), Rab7 (late endosome), Rab11 (recycling endosome), LAMP1 (lysosome), and GM130 (Golgi). PNS=post-nuclear supernatant. **(B)** Quantification of NHE6 protein levels in endosome fractions in HAP1 GGA1-KO line 1 and WT cells. (mean±SEM; n=2; *p<0.05, unpaired *t* test). **(C)** Western blot of NHE6 protein in late endosome fractionation in HAP1 GGA1 KO line 2 and WT cells using the following markers: Rab5 (early endosome), Rab7 (late endosome), Rab11 (recycling endosome), LAMP1 (lysosome), and GM130 (Golgi). PNS=post-nuclear supernatant. **(D)** Quantification of NHE6 in late endosome fractionation in HAP1 GGA1KO line 2 and WT cells. (mean±SEM; n=2; unpaired *t* test).

### Loss of GGA1 disrupts NHE6 cellular distribution

We performed lysosome compartment fractionation in HAP1 cells to determine how GGA1 affects the distribution of NHE6 in lysosomes. As shown from Figure 7A, NHE6 is found in the fractionated lysosome compartment, using LAMP1 as a lysosome marker. NHE6 was significantly elevated in the fractionated lysosome compartment in GGA1 KO (Figure 7B). This implies that loss of GGA1 promotes mislocalization of NHE6 to lysosomes.

**Figure 7.**
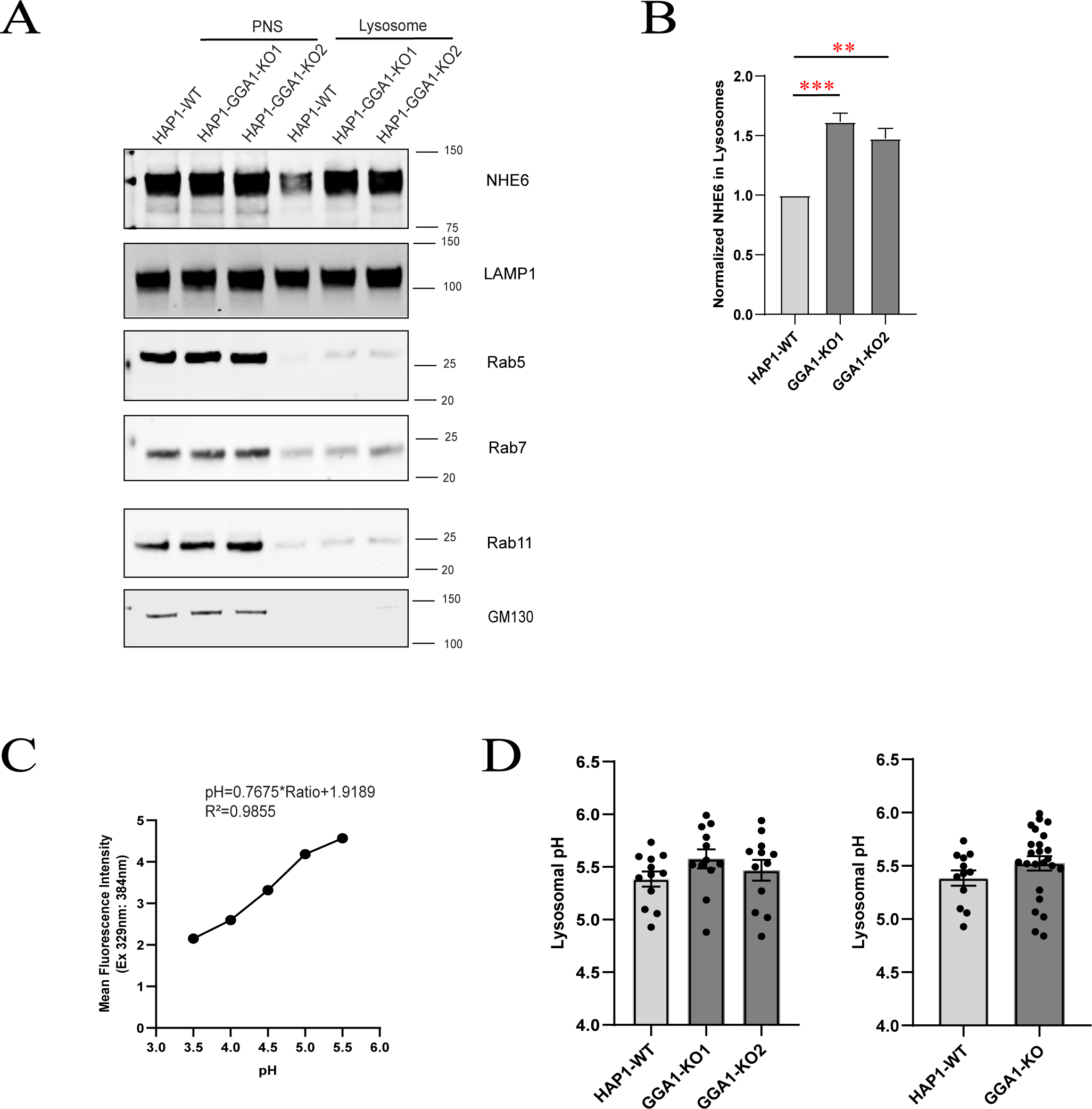
Loss of GGA1 leads to greater NHE6 in lysosomes, but does not affect lysosome pH. **(A)** Western blot of NHE6 protein in lysosome fractionation in HAP1 GGA1 knockout (KO) and wildtype (WT) cells using the following markers: LAMP1 (lysosome), Rab5 (early endosome), Rab7 (late endosome), Rab11 (recycling endosome), GM130 (Golgi). PNS=post-nuclear supernatant. **(B)** Quantification of NHE6 in lysosome fractionation in HAP1 GGA1 KO and WT cells. (mean±SEM; n=3; **p<0.01, ***p<0.001, unpaired *t* test). **(C)** pH calibration curve graph for lysosome pH measurement by LysoSensor™ Yellow/Blue DND-160 after 1 min incubation. **(D)** Quantification of lysosomel pH in HAP1 GGA1 KO and WT cells. (mean±SEM; n=12 replicates; unpaired *t* test)

Since loss of NHE6 leads to hyper-acidification of the lysosome compartment (8), we wondered whether enriched NHE6 localization in lysosomes in GGA1 KO lines would lead to alkalinization of lysosome pH. We measured lysosome pH using the unconjugated pH-dependent fluorescent probe LysoSensor DND-160. Considering the risk of lysosome alkalization as an effect of LysoSensor in long time treatment, we used a short 1min treatment time at the concentration of 1uM according to the instructions described in experimental procedures. We calculated lysosome pH using the equation derived from a standard curve and converting the mean intensity to mean lysosomal pH (Figure 7C). There were no statistically significant differences in intra-luminal lysosomal pH between GGA1 KO and control lines (Figure 7D). From these findings we conclude that loss of GGA1 leads to greater distribution of NHE6 to lysosomes while lysosome pH is unaffected.

We then performed Golgi fractionation in GGA1 KO HAP1 cells to determine whether loss of GGA1 alters NHE6 Golgi localization. We did not detect any statistically-significant differences in the amount of NHE6 in the Golgi fraction between GGA1 KO and control cells (Figure 8B). We then measured luminal TGN pH using a ratiometric TGN38-pHluorin construct in GGA1 KO HAP1 cells (64–66). We first confirmed that TGN38-pHluorin is properly trafficked to Golgi compartments as it colocalizes with the cis-Golgi, (GM130, Figure S6A) and trans-Golgi network (TGN38, Figure S6B) markers. Loss of GGA1 caused a significant increase in trans-Golgi network pH compared to the control HAP1 line (Figure 8D). These findings suggest that loss of GGA1 leads to alkalinization of the trans-Golgi network, possibly mediated by NHE6 activity.

**Figure 8.**
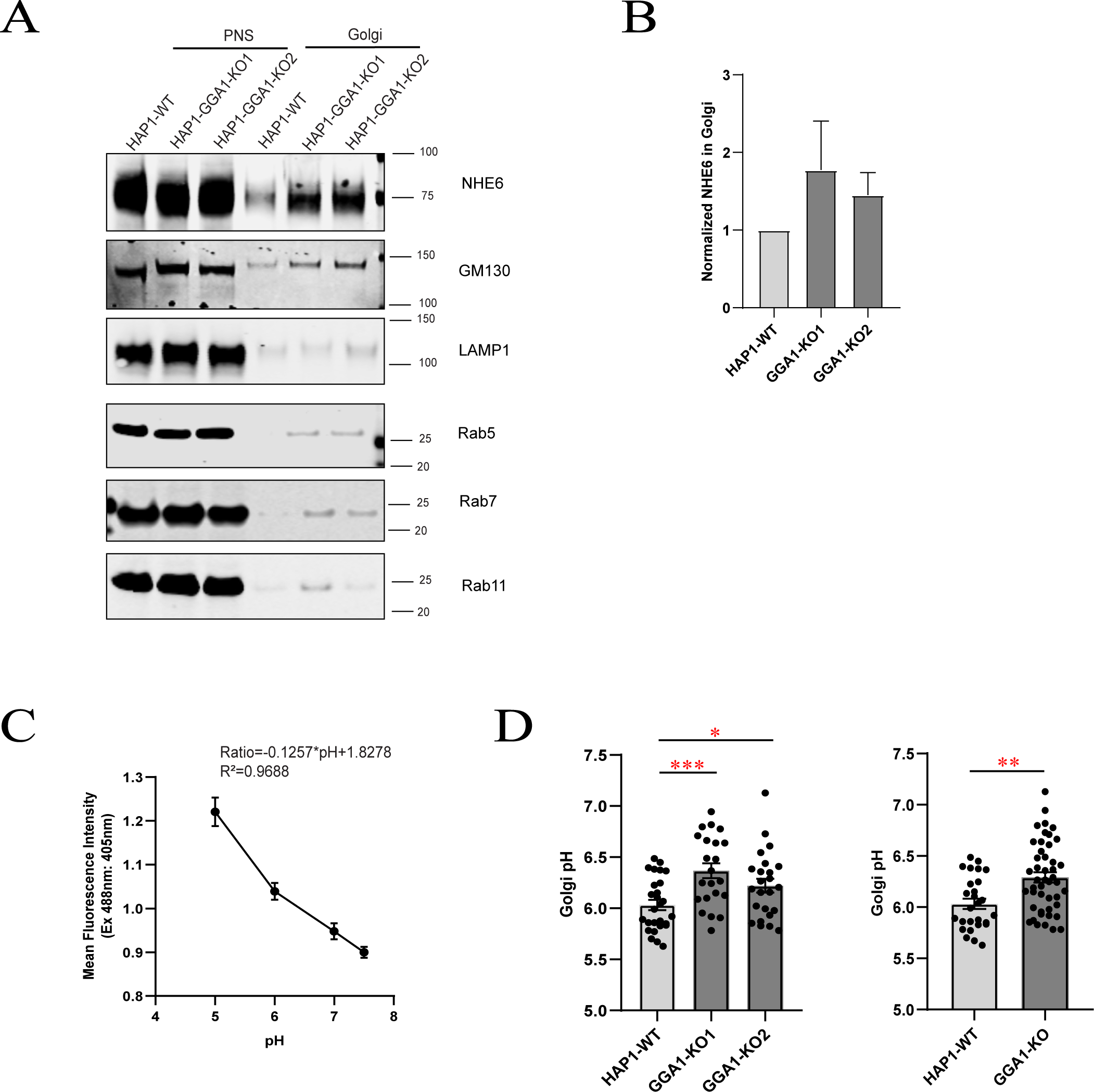
Loss of GGA1 does not alter NHE6 Golgi distribution, but alkalinizes Golgi pH. **(A)** Western blot of NHE6 protein in Golgi fractionation from HAP1 GGA1 knockout (KO) and wildtype (WT) cells using the following markers: GM130 (Golgi), LAMP1 (lysosome), Rab5 (early endosome), Rab7 (late endosome), and Rab11 (recycling endosome). PNS=post-nuclear supernatant. **(B)** Quantification of NHE6 in Golgi fractionation in HAP1 GGA1 KO and WT cells. (mean±SEM; n=3; unpaired *t* test). **(C)** Graph of the Golgi pH calibration curve in HAP1 GGA1 KO and WT cells using TGN38-pHluorin construct. **(D)** Quantification of Golgi pH in HAP1 GGA1 KO and WT cells. (mean±SEM; WT n=27 GGA1 KO1 n=23, GGA1 KO2 n=24 cells; *p<0.05 and *** p<0.001, unpaired *t* test).

Finally, we examined whether loss of GGA1 alters NHE6 distribution to the cell surface, as knocking down GGA1-3 by siRNA in HeLa cells increases extracellular secretion of the lysosome enzyme cathepsin D (38). As shown from Figure 9, GGA1 KO cells exhibited significantly more NHE6 on the cell surface compared to control cells. Therefore, loss of GGA1 leads to altered trafficking of NHE6 to the plasma membrane. This result suggests that the interaction of NHE6 with GGA1 may promote NHE6 endosome localization and loss of GGA1 enables NHE6 exocytosis from Golgi to the cell surface. Overall, in the absence of GGA1, we observe mislocalization of NHE6 in the cell: we observe physiologically relevant NHE6 elevations in Golgi, reduced levels trafficking to endosome, elevations in lysosome, as well as elevations on the cell surface.

**Figure 9.**
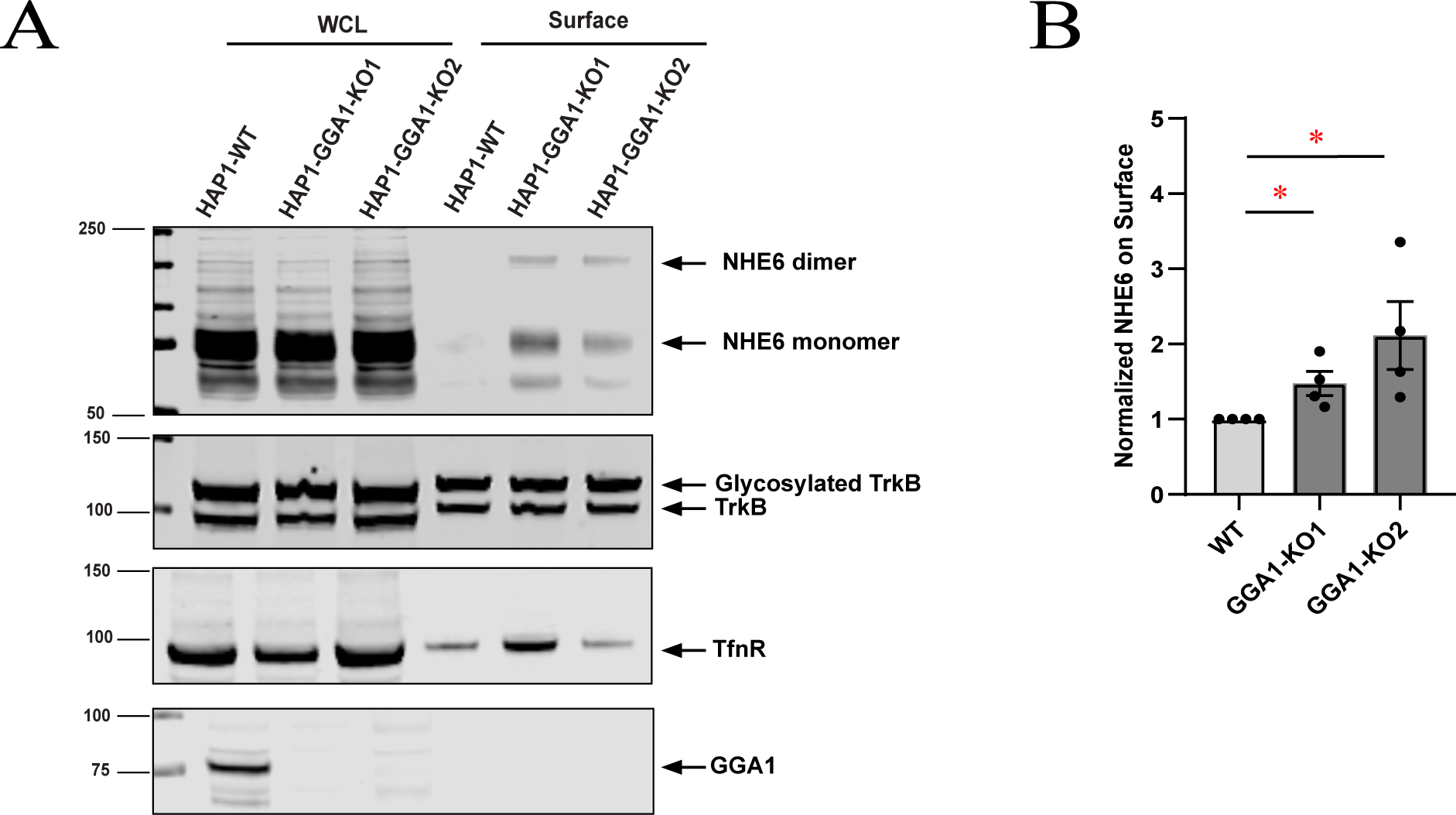
Loss of GGA1 leads to more NHE6 on plasma membrane. **(A)** Whole cell lysate (WCL) and biotinylated surface were separated by Western blot and detected by immunoblotting with anti-NHE6, transferrin (TfR) and TrkB antibodies. **(B)** Quantification of NHE6 on plasma membrane in HAP1 GGA1 KO and WT cells. (mean±SEM; n=4; *p<0.05, unpaired *t* test).

## DISCUSSION

NHE6 functions in regulating of intra-endosomal pH and endosome maturation (4,8), yet the mechanism of this process is incompletely understood. In this study, we identify new NHE6-interaction partners – GGA1 and GGA3. We demonstrate an interaction between the C-terminus of NHE6 and the GGA1 protein by two-hybrid screening. A strong result of 13 among 123 captured clones in the yeast-two hybrid encoded the GGA1 protein, indicating the C-terminus of NHE6 is sufficient for binding with GGA1 in yeast cells. The chimeric NHE1/NHE6 experiment further indicates that the NHE6 C-terminus is necessary and sufficient for the interaction with GGA1. We further demonstrate that this interaction occurs in endogenously expressed GGA1 and NHE6 in mammalian cells, and mouse brain tissue.

We further demonstrate that GGA1 predominantly interacts with organellar NHEs (e.g. NHE6, NHE7 and NHE9), as opposed to cell surface localized NHEs (e.g. NHE1 and NHE5). Additionally, we show that GGA3 interacts with organellar NHE6 and NHE9. Taken together, these findings indicate that organellar NHE family members, that have high C-terminus similarity, are new interaction partners with GGA family members. Overall, these data suggest that GGA1 may function in transporting NHEs between the Golgi and endocytic pathway.

We characterized the GGA1 domains responsible for binding with NHE6. The GGA1 domains VHS, GAT, and GAE regions all interact with NHE6, with the GAE domain showing the strongest interaction with NHE6. Further experiments will be required to address the underlying mechanism regarding the strong interaction of GAE domain with NHE6. Although the GAE domain strongly interacts with NHE6, the “Hinge linker and GAE” domain reduces this interaction., This finding suggests that the Hinge linker domain functions as a dominant negative. In the Hinge linker domain, the AC-LL motif functions as an autoinhibitor by competing to bind the ligand-banding site in VHS domain (37,55). In this study, it is possible the AC-LL motif in GGA1 Hinge linker domain (^357^DDELM^361^) may play an autoinhibition role on NHE6 versus GGA1 VHS domain interaction. NHE6 plasmids used in this study contain the NHE6.2 isoform, which share the same C terminus sequence with the canonical NHE6.1. The major difference between these isoforms is that NHE6.2 has a shorter N terminus by 32 amino acid residues, which does not appear to affect the interaction between NHE6 and GGA1/3.

Using two HAP1 GGA1 KO cell lines, we find loss of GGA1 alters the distribution of NHE6. In GGA1 KO cells, NHE6 protein was less likely to be found in early endosomes and more likely to be found in lysosomes and at the plasma membrane. This is consistent with loss of GGAs leading to greater extracellular secretion of lysosomal enzymes via the plasma membrane (34). Given the central role of NHE6 is to alkalinize endolysosomal compartments (4,8), we examined whether NHE6 mislocalization in GGA1 KO cells disrupts organellar pH. Despite enriched NHE6 in lysosomes in GGA1 KO cells, there were no significant differences in lysosomal pH. However, Golgi pH was significantly increased (i.e. more alkaline) in GGA1 KO cells compared to WT cells. Taken together, these findings expand our understanding of how GGA1 regulates NHE6 trafficking and the functional consequences of disrupting this interaction. Loss of GGA1 alters the distribution of NHE6. In HAP1 GGA1 KO cell lines, NHE6 is less likely to be found in early endosomes, whereas it is enriched in lysosomes and at the cell surface. Previous reports have indicated that minimal NHE6 is localized to lysosomes (12,61). Despite enriched NHE6 trafficking to lysosomes, we did not detect a difference in pH in the lysosomal lumen. Therefore, we posit that the lysosome-associated NHE6 is not a key regulator for lysosomal pH and/or is non-functional. Interestingly, Golgi fractionation showed that more NHE6 accumulated in the Golgi in the absence of GGA1, at the same time, an alkalinized Golgi lumen was detected consistent with NHE6 retention in the Golgi.

Loss of NHE6 leads to Christianson syndrome, a neurogenetic disorder associated with neurodegenerative features including progressive cerebellar atrophy, motor decline, and tau depositions neurodegeneration in a condition called Christianson Syndrome (CS), which appears to be due in part to endolysosome dysfunction (8,10,67,68). An important question related to CS is the extent to which underlying pathogenic mechanisms are shared with more common neurodegenerative conditions such as Alzheimer’s Disease (AD). The interaction between NHE6 and GGA1 supports these ideas of overlap; for example, GGA1 regulates the levels of β-site amyloid precursor protein (APP)-cleaving enzyme 1 (BACE1) (39,69), and also modulates the processing of amyloid precursor protein (APP) to amyloid-β (Aβ) (70–72). So, the interaction between GGA1 and NHE6 may promote less aggregation of Aβ in neurodegenerative defects like AD. This study introduces a new NHE6-interacting protein, and also opens up new directions in the exploration of overlap between neurodegenerative mechanisms in CS and in other neurodegenerative diseases such as AD.

Our study establishes a novel NHE6 binding partner – GGA1 – using different models and multiple techniques including yeast two-hybrid, IP, subcellular fractionation, super-resolution and expansion microscopy. GGA1 domains VHS, GAT, and GAE are all able to bind NHE6. We extend our investigation to identify which NHEs interact with GGA1. We find that GGA1 interacts with other organellar NHEs like NHE6 (e.g. NHE7 and NHE9), but not plasma membrane NHEs (e.g. NHE1 and NHE5). We also find that another GGA family member– GGA3 – interacts with the organellar NHEs NHE6 and NHE9. It remains unknown whether plasma membrane NHEs are more likely to bind other GGA family members. This work highlights the relationship between GGAs and NHEs. Using super-resolution and expansion microscopy we visualize the distribution of NHE6-GGA1 colocalization in neurons *in vitro*. These experiments demonstrate that NHE6-GGA1 colocalization occurs predominantly in the perinuclear region in the Golgi complex and elsewhere, presumably endosomes. We discover that loss of GGA1 alkalinizes the trans-Golgi network. We believe NHE6 is involved in this finding given (1) GGA1’s role in NHE6 trafficking and (2) NHE6’s role in regulating pH. It is unlikely this is due to greater NHE6 levels in the TGN as there were no differences in NHE6 protein levels in our Golgi fractionation experiment. Therefore, we suspect enhanced NHE6 exchanger activity mediates the higher TGN pH in GGA1 KO cells. Abnormal Golgi pH impairs glycosylation and protein trafficking and is associated with a range of human diseases (73,74). Notably, loss of the Angelman syndrome protein Ube3a leads to an overlapping similar Golgi pH phenotype as loss of GGA1, as the Golgi complex becomes hyper-alkalinized (75). A limitation of our study is that LysoSensor DND-160 does not directly measure lysosome pH, but rather the pH of acidic organelles. While LysoSensor is most likely to measure the luminal pH of acidic lysosomes, it may also include highly acidic endosomes. Thus, our finding that loss of GGA1 enhances NHE6 localization to lysosomes, but does not affect lysosome pH suggests that NHE6 exchanger function is less active in lysosomes.

In summary, our study identifies GGA1 as a new binding partner with NHE6 and establishes a link between NHEs and GGAs. We find that NHE6 and GGA1 localize throughout the cell including endosomes and the Golgi complex. Loss of GGA1 leads to NHE6 mis-localization as its expression is increased in lysosomes and the plasma membrane, but decreased in endosomes. Functionally, loss of GGA1 alkalinizes TGN pH that is likely mediated by NHE6.

## EXPERIMENTAL PROCEDURES

### Yeast two-hybrid screening and assay

The coding sequence for amino acids Leu^537^-Asp^630^ of the mouse NHE6 protein (GenBank accession number gi: 120577706) was PCR-amplified and cloned into pB27 as a C-terminal fusion to LexA (N-LexA-NHE6-C). The peptide sequence corresponded to: LHIRVGVDSDQEHLGVPDNERRTTKAESAWLFRMWYNFDHNYLKPLLTHSGPPLTTTLPACCGPIARCLTSPQAYENQEQLKDDDSDLILNDGDISLTYGDSTVNTESATASAPRRFMGNSSEDALDRELTFGDHELVIRGTRLVLPMDDSEPALNSLGDTRHSPA. The construct was checked by sequencing the entire insert and used as a bait to screen a random-primed rat hippocampus cDNA library constructed into pP6. pB27 and pP6 derive from the original pBTM116 (76) and pGADGH (77) plasmids, respectively.

53.8 million clones (5.4-fold the complexity of the library) were screened using a mating approach with Y187 (matα) and L40ΔGal4 (mata) yeast strains as previously described (78). 123 His+ colonies were selected on a medium lacking tryptophan, leucine and histidine, and supplemented with 0.5 mM 3-aminotriazole to handle bait autoactivation. The prey fragments of the positive clones were amplified by PCR and sequenced at their 5’ and 3’ junctions. The resulting sequences were used to identify the corresponding interacting proteins in the GenBank database (NCBI) using a fully automated procedure. A confidence score (PBS, for Predicted Biological Score) was attributed to each interaction as previously described (79).

### Further description of the confidence score

The Predicted Biological Score (PBS) relies on two different levels of analysis. Firstly, a local score takes into account the redundancy and independency of prey fragments, as well as the distribution of reading frames and stop codons in overlapping fragments. Secondly, a global score takes into account the interactions found in all the screens performed using the same library. This global score represents the probability of an interaction being nonspecific. For practical use, the scores were divided into four categories, from A (highest confidence) to D (lowest confidence). A fifth category (E) specifically flags interactions involving highly connected prey domains previously found several times in screens performed on libraries derived from the same organism. Finally, several of these highly connected domains have been confirmed as false-positive of the technique and are now tagged as F. The PBS scores have been shown to positively correlate with the biological significance of interactions (80,81).

### Cell culture and reagents

HEK293T cells were cultured in Dulbecco’s modified Eagle’s medium (DMEM) supplied with 10% fetal bovine serum (FBS), 1% Antibiotic-Antimycotic and 1% GlutaMAX™, SH-SY5Y neuroblastoma cells were cultured in DMEM/F12 medium supplied with 10% fetal bovine serum, 1% Antibiotic-Antimycotic, HAP1 WT and GGA1-KO cells were cultured in Iscove’s Modified Dulbecco’s Medium (IMDM) supplied with 10% FBS and 1% Antibiotic-Antimycotic at 37°C and 5% CO2. All cell culture media and reagents were obtained from Invitrogen (Carlsbad, CA). Rabbit polyclonal anti-NHE6 antibody was customized made against isoform-specific sequences GDHELVIRGTRLVLPMDDSE (aa636-655) of the C terminus of NHE6.0 (4). Antisera were collected and affinity-purified (4). HA antibody (Cell signaling, 3724s) and Santa Cruz (sc-7392), GFP antibody (Cell signaling, 2956s), GGA1 antibodies (Novus Biologicals, H00026088-M01 (3F11)) and Pierce (PA5-12130). GGA3 antibody (BD Biosciences, 612310), GM130 (BD Biosciences, 610822), LAMP1 (Abcam, ab25630), TrkB (BD Biosciences, 610101). Rab5 (Cell Signaling, 3547s), Rab7 (Abcam, ab137029) and Rab11(BD Biosciences, 610656). Transferrin receptor (Invitrogen, 13-6890), NHE5 (PA5-37222), NHE7 (PA5-75424) and NHE9 (Morrow lab customized made through Covance, epitope located within the C-terminal tail of mouse Slc9a9: SPSPSSPTTKLALDQKSSGKC). Flag M2 (Sigma, F1804). a-Tubulin (Sigma, T6074), c-Myc antibody (Santa Cruz, sc-40). Protein G agarose beads was purchased from Santa Cruz (sc-2002) and Dynabeads^TM^ Protein G from Thermo Fisher Scientific, 10003D. Lysosome enrichment kit (Thermo Fisher Scientific, 89839) and Golgi isolation kit (Sigma, GL0010). Pierce™ cell surface protein biotinylation and isolation kit (Thermo Fisher Scientific, A44390). LysoSensorTM Yellow/Blue DND160 (Invitrogen**™,** L7545). Sytox^TM^ Green Nucleic acid stain (S7020, Thermo Fisher Scientific).

### GGA1 Knockout cell lines

Two GGA1 knockout cell lines (KO1 and KO2) were customized generated by Horizon Discovery. Detailed information is shown in Table S2.

### Plasmids constructs, PCR-mediated mutagenesis and transfection

pCMV-mGGA1 which was originally from Open Biosystems, GGA1 full length and domains were amplified from pCMV-mGGA1 and then cloned into mammalian expression vector pcDNA3.1/CT-GFP-TOPO (Invitrogen). PCR-mediated GGA1-Hinge domain (299-505aa) mutagenesis was carried out according to the methods provided by Quickchange II Site-Directed Mutagenesis Kit. Myc-GGA1 was amplified from pCDNA3.1-mGGA1 and double digested by restriction enzymes of EcoRI +XhoI and then ligated to Clontech c-Myc-vector.

pmNHE9 was originally from Open Biosystems, and then cloned into mammalian expression vector pcDNA3.1/CT-GFP-TOPO (Invitrogen). HA tagged hNHE6.2-FL and Cytoplasmic domain (CD) were cloned by GeneCopoeia. hNHE1-HA (3X on c-terminal) (pYN4+, #78715) was originally from Addgene with D720G mutation, mutagenesis was performed to correct this mutation back. hNHE5 (#132163), hNHE7 (#132187), which was originally obtained from Addgene, together with mNHE9-GFP, cloned into HA vector using HA tagged hNHE6.2-FL as template and using in-fusion snap method (Takara, In-Fusion® Snap Assembly Master Mix, 638947) to replace hNHE6 with hNHE5, hNHE7 or mNHE9.

In-fusion snap method was also used for chimeric NHE6/NHE1 plasmids construction. Snap NHE1N/NHE6C-HA was constructed by replacing hNHE6 N-terminus (1-504aa) with hNHE1.1 N-terminus (1-499aa). Snap NHE6N/NHE1C-HA was constructed by replacing hNHE6 C-terminus (505-669aa) with hNHE1.1 C-terminus (500-815aa).

Transfection of cells with various mammalian expression constructs by Lipofectamine 2000 (Invitrogen, Carsbad, CA, USA) was according to the methods provided by manufacturer’s specification.

All constructs were finally verified by DNA sequencing by Genewiz or PlasmidSaurus. All primers were listed in Table S3 and 4.

### Western blotting and co-immunoprecipitation

Cells/Tissues were lysed in buffer containing: 50 mM Tris-HCl, pH 7.8, 137 mM NaCl, 1 mM NaF, 1 mM NaVO3, 1% Triton X-100, 0.2% Sarkosyl, 1 mM dithiothreitol (DTT), and 10% glycerol or buffer containing: 50mM Tris-HCl pH7.9, 100mM NaCl, 20% glycerol and 0.1% NP-40 or RIPA buffer supplemented with protease inhibitor cocktail and phosphatase inhibitor. All cells were lysed for 30 min on ice and were centrifuged at 13,200 rpm for 15 min at 4°C to remove cell debris. Protein concentration was measured by BCA assay using the Pierce BCA Kit (Thermo Fisher Scientific 23225). For immunoprecipitation, antibody was conjugated to Dynabeads Protein G (Thermo Fisher Scientific 10003D) at room temperature for 2 hr. Cell lysates were then incubated with antibody-conjugated beads overnight (O/N) at 4 ◦C, or cell lysates were incubated with antibody for 2hr then conjugate Protein G plus-agarose (Santa Cruz, sc-2002) (O/N) at 4 ◦C. The following day, the beads were gently pelleted, cell lysates were removed, and the beads were washed three times with PBS with 0.02% TWEEN 20 wash buffer. Pelleted beads were then boiled in sample buffer at 95 ◦C for 5 min before loading onto 4–12% SDS-PAGE gels (Novex #NP0321Box). Following separation of proteins by electrophoresis, gels were transferred to nitrocellulose membranes (Novex #LC2000). Western blots were performed using standard procedures (5,82) and were analyzed with the Li-CoR Odyssey Imaging System.

### Primary cultured neurons immunocytochemistry and SIM Deltavision imaging

Primary neurons were dissociated and as described before (4). For hippocampal cultures, wild-type neurons were firstly washed three times with 1× PBS, then fixed with 4% paraformaldehyde for 10 min at room temperature and permeabilized for 10 min in PBS containing 0.25% Triton X-100. Non-specific binding was blocked by incubation with 10% normal goat serum (Jackson ImmunoResearch #005-000-121) in PBS containing 0.1% TWEEN 20 (PBST) for 1hr. Cells were then incubated overnight at 4 ◦C with primary antibody diluted in PBST containing 2% normal donkey serum, washed 3 × 5 min with PBST, and incubated for 1 h at room temperature with secondary antibody diluted as for primary antibody. Nuclei were counterstained with Hoechst (1:1600 working dilution of 10 mg/mL stock; Invitrogen #33342). Cells were then washed 3 times with PBST and mounted on slides with Fluoromount-G (SouthernBiotech #0100-01) or ProLong™ Glass Antifade Mountant (Invitrogen P36984). Images were captured with SIM deltavision or FV3000 Confocal or Zeiss LSM800 microscope.

SIM images were collected using a DeltaVision OMX SR microscope. Z-series images were collected using a 60× oil objective (refractive index immersion oil 1.516) and structured illumination (SI) light path under sequential acquisition mode. Z-series images were processed by performing OMX SI reconstruction, alignment, and maximal projection sequentially using softWoRx software. Images were analyzed using ImageJ software (NIH).

### Co-localization of GGA1 and NHE6 by 5x expansion microscopy method

#### Cell culture expansion

Rat hippocampal neurons were cultured from wildtype rat pups (p0-p2) in neurobasal media supplemented with B27 and Glutamax 1%. On DIV 14 rat hippocampal neurons were fixed with 4% PFA for 30 mins and followed by post-fixation and first round of gelation for hydrogel expansion (5x) as per the protocol mentioned in M’Saad O. and Bewersdorf J, 2020 (57).

#### Antibody labelling

For GGA1 and NHE6 co-localization with respect to Golgi/Endosomes in 5x expanded cells in hydrogels were labelled with a cocktail of primary antibodies (1:500) of mouse GGA1 (H00026088-M01, Novus Biologicals), rabbit NHE6 (Covance), and guinea pig Giantin (263 004, Synaptic Systems)/guinea pig EEA1 (237 105, Synaptic Systems) were incubated in TBS-tween (927-65001, LI-COR Biosciences) for 24 h on a rocking platform at RT. The expanded gels were later washed 3 times with TBS-tween 15 mins each followed by a secondary antibody labelling. Later incubated in the cocktail of secondary antibodies (1:500) Goat anti-guinea pig Alexa488, Donkey anti-mouse CF568, and Goat anti-rabbit Alexa647 in TBS-tween for 16-20 h, followed by three 15 min washes with PBS-tween. The nucleus was stained with 1:1500 Sytox^TM^ Blue Nulceic acid stain (S11348, Thermo Fisher Scientific) in 1x PBS for 30 mins followed by three 15 min washes in 1x PBS. The water was changed three times every 30 mins and incubated in water O/N at RT.

#### Imaging

On the day of imaging the gels were mounted glass bottomed Mattek dishes and sealed with dental glue as described in M’Saad O. and Bewersdorf J, 2020 (57). For cells expanded using hydrogels, z-stacks were acquired using Olympus FV3000 microscope. Images were collected with 1024×1024 pixel resolution using 60x water objective. Voxel size for the acquired images were ∼114×114×63 uM.

#### Analysis

For GGA1 and NHE6 co-localization with respect to Golgi/Endosomes in 5x expanded cells in hydrogels, IMARIS software (version 10.0.0.1) was used for analysis. Nucleus and Giantin/EEA1 were analyzed using surfaces visualization in the surpass tree of IMARIS software. GGA1 and NHE6 localizations were analyzed using spots visualization. The overall and shortest distance statistics were generated and extracted from the software to plot using GraphPad Prism version 7. The statistics were generated from the perspective of both GGA1 and NHE6 and plotted in the same graph. For measuring expansion factor contour was drawn around the nucleus (max projection) using the magic wand tool in FIJI. The area of the nucleus was measured using measure plugin from FIJI. The expansion factor, (EF) was calculated by measuring square root of area of expanded neurons, (a2) by area of unexpanded neurons, (a1) from the max projection. EF = √(a2/a1).

### Early and late endosome subcellular fractionation

Early and Late endosomal fractions were prepared as described (62,63). 6∼8×10^7^ cells used in this study were placed on ice, washed, scraped into microfuge tubes and homogenized (Sucrose 250mM, Imidazole (pH7.4) 3mM, EDTA 1mM, Cycloheximide 0.03mM with protease inhibitors and phosphatase inhibitors added), and then a post-nuclear supernatant (PNS) was prepared. The PNS was adjusted to 40.6% sucrose (refractometry was used to accurately measure the % of sucrose) in 3 mM imidazole, pH 7.4, loaded at the bottom of an ultra-thin centrifuge tube, and overload sequentially with 1.5 volumes of 35% and 1 volume of 25% sucrose solutions in 3 mM imidazole, pH 7.4, and then homogenization buffer (HB; 250 mM sucrose in 3 mM imidazole, pH 7.4). The gradient was centrifuged for 3hrs at 41,600 rpm using a SW55Ti rotor (Table S5). Early and late endosomal fractions were collected at the 35/25% and 25%/HB interfaces, respectively. Remove the bands by needle puncture on the tube wall to get EE and LE fractionation. BCA assay was applied, and equal amount of protein was loaded for WB assay and different fractionation makers were used to monitor the purity of EE/LE fractionation.

### Lysosome subcellular fractionation

Lysosome Enrichment Kit (Thermo Fisher Scientific 89839) was used for tissue and cultured Cells lysosome subcellular fractionation according to manual’s instructions. TLA-110 rotor was used for ultracentrifuge at 145,000 × g for 2 hours (Table S5) at 4°C. Refractometry was used to accurately measure the % of sucrose.

The transparent lysosome band is on the top 2mL of the gradient and under thick lipid. Carefully remove the lysosome band and save on ice. BCA assay was applied, and equal amount of protein was loaded for WB assay and different fractionation makers were tested to monitor the purity of lysosome fractionation.

### Golgi subcellular fractionation

Golgi isolation Kit (Sigma GL0010) was used with some modifications for cultured Cells Golgi subcellular fractionation. Cells were homogenized in 0.25 M Sucrose Solution and centrifuge to get supernatant. Adjust the sucrose concentration in the sample (supernatant) to 1.25 M by adding the volume of 2.3 M Sucrose Solution according to formula provided by kit. Build a discontinuous gradient and the order of sucrose gradient fractions in the ultracentrifuge tube (from bottom to top) should be: 1.84 M Sucrose Solution; Sample (sucrose concentration adjusted to 1.25 M); 1.1 M Sucrose Solution and 0.25 M Sucrose Solution. The gradient was centrifuged for 3hrs at 120,000 g using a SW55Ti rotor for 3 hours (Table S5) at 2–8 °C and then withdraw the Golgi enriched fraction from the 1.1 M/0.25 M sucrose interphase. BCA assay was applied, and equal amount of protein was loaded for WB assay and different fractionation makers were tested to monitor the purity of Golgi fractionation.

### LysoSensor^TM^ Yellow/Blue DND160 lysosomal pH measurement

HAP1 WT and GGA1 KO cells were incubated at 37°C in pre-warmed, IMDM culture medium containing 1 μM of LysoSensor™ Yellow/Blue DND-160 for 1min. For those wells prepared for pH calibration curve, rinse cells 2 times with 1XPBS and once with pH calibration curve buffer (3.5, 4, 4.5, 5, 5.5) and then Incubate for 10 min in 100 μL of respective pH calibration curve buffer prepared for reading in SpectraMax® M5 Microplate Reader. Light emitted at 440 and 540nm in response to excitation at 329 and 384nm was measured. The ratio of light emitted at 340 and 380nm excitation was plotted against the pH values and the pH calibration curve for the fluorescence probe was generated from the plot using Microsoft Excel. Calculate the pH using the equation from pH calibration curve (64).

### Golgi compartment pH measurement

HAP1 WT and GGA1 KO cells were seeded to 35mm glass bottom dishes for next day transiently transfected with TGN38-pHluorin plasmid for 24hrs before taking images. For standard curve making, rinse cells once with the most alkalized pH standard buffer for imaging, find appropriate region of Golgi pHluorin-expressing cells for imaging based on excitations at 405 nm and 488 nm and both emitted at 530 nm. Then rinse cells once with 1X PBS followed by rinses twice with the next pH calibration curve buffer from alkalized to acidic. Collect images for each pH calibration curve. During image taking, maintain cells on a heated stage held at 37°C in a CO2 chamber. For sample images taking, rinse cells one time with PBS and keep cells in phenol free medium during images taking. Image J was then used to analyze data and regions of interest within cells containing pHluorin-labeled Golgi were selected. Export florescence intensity measurement data to excel and calculate the fluorescence intensity ratio of intensity of two excitation wavelengths for each region of interest. Calculate the Golgi pH using the equation generated from making pH calibration curve (64).

### Cell surface protein biotinylation and isolation

HAP1 WT and GGA1 KO cell surface protein biotinylation and isolation were performed by following the Pierce cell surface protein biotinylation and isolation kit’s manual instruction (Thermo Fisher Scientific A44390). In brief, cells were washed with PBS before labelling with Sulfo-NHS-SS-Biotin for 10min at RT. Then the label was removed, washed and cells were harvested for lysis. An equal amount of supernatant fraction from lysed cells was incubated with NeutrAvidin Agarose slurry for 30min at RT. After 4 times of washing, biotinylated protein was eluted by incubating with elution buffer for 30min at RT. The biotin-labeled surface proteins were separated by SDS-PAGE and analyzed by immunoblotting. The intensity of the immunoreactive bands was quantified using Li-COR analysis software.

## Acknowledgments

This work was supported by NIH NINDS grants (R01NS113141, R01NS121618) to E.M.M. The authors wish to acknowledge Dr. Judy S. Liu for her scientific and technical discussion. The authors also acknowledge Dr. Terry E. Machen from University of California-Berkeley, Berkeley, CA for providing TGN38-pHluorin plasmid. The authors also acknowledge Dr. Mattew F. Pescosolido for assistance in manuscript preparation. The authors thank Dr. Walter Atwood, Bethany O’Hara and Genomics Core Dr. Christoph Schorl from Brown-Department of Molecular Biology, Cell Biology and Biochemistry for their assistance on the usage of ultracentrifuge machines.

## Author contributions

Conceptualization L.M. and E.M.M.; Data Curation and Visualization L.M., Q.O., M.S.; 5x expansion microscope imaging and analysis by R.K.K.; Experimental Design and Data Analysis L.M. and E.M.M.; Formal Analysis and Writing – Original Draft, L.M. and E.M.M.; Writing – Review & Editing, L.M. and E.M.M; Supervision, Project Administration, and Funding Acquisition, E.M.M.

## SUPPLEMENTAL FIGURE LEGENDS

**Figure S1.**
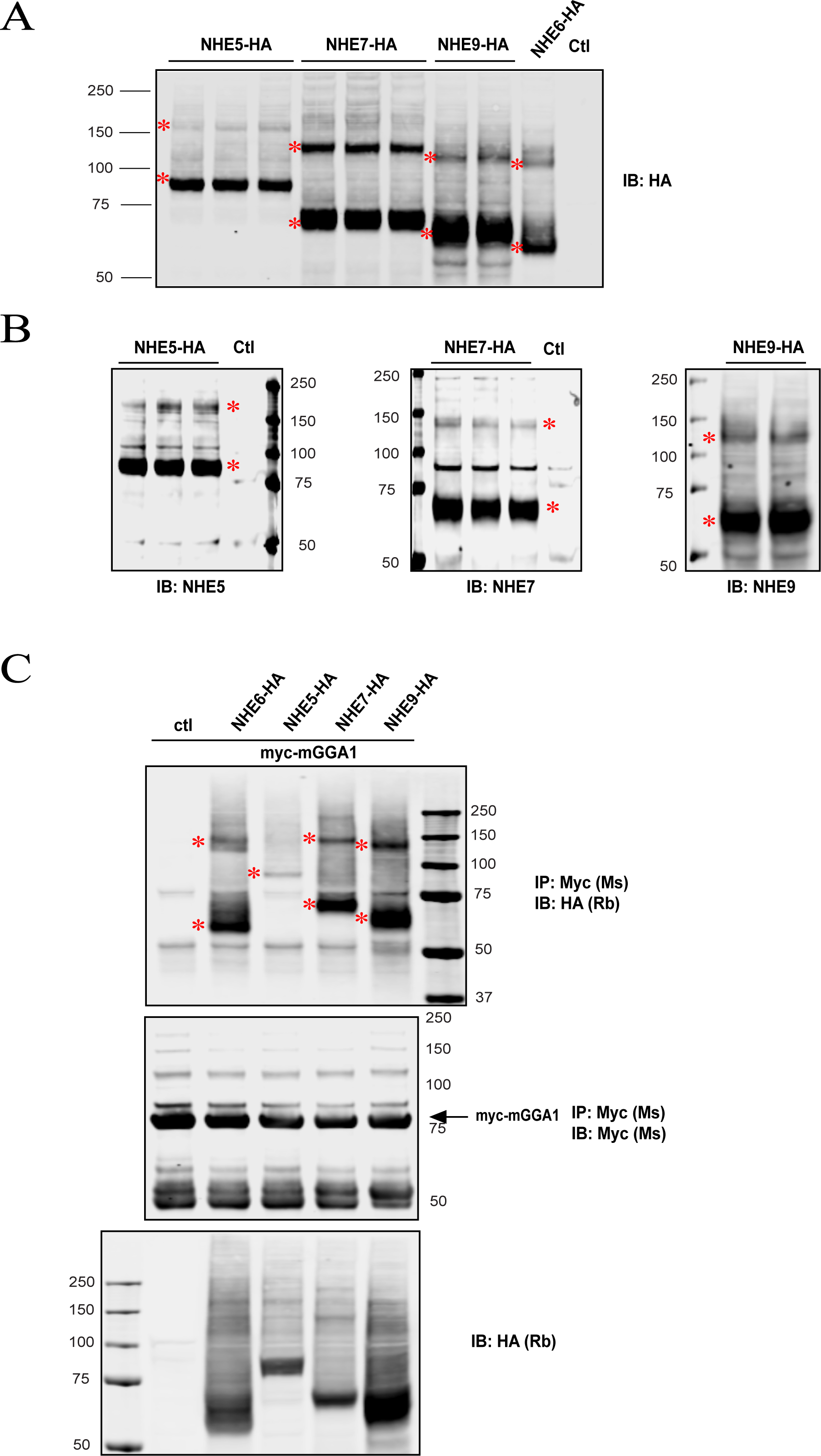
GGA1 interaction with NHEs. **(A)** Cell lysates from HEK293T cells expressing HA-tagged NHE5, NHE7, and NHE9 were subjected to western blot analysis with anti-HA to detect expression of NHE5, NHE7, and NHE9 constructs. NHE6-HA was used as a control. **(B)** Cell lysates from HEK293T cells expressing HA-tagged NHE5, NHE7, and NHE9 were subjected to western blot analysis with anti-NHE5, NHE7, and NHE9 to detect expression of NHE5, NHE7, and NHE9 constructs. **(C)** Cell lysates from HEK293T cells expressing c-Myc-GGA1 with HA-tagged NHE5, NHE7, NHE9, and NHE6 were immunoprecipitated with an anti-c-Myc antibody. The precipitates were probed with HA and c-Myc antibodies. Western blot analysis was performed with anti-HA to detect the expression of NHEs.

**Figure S2.**
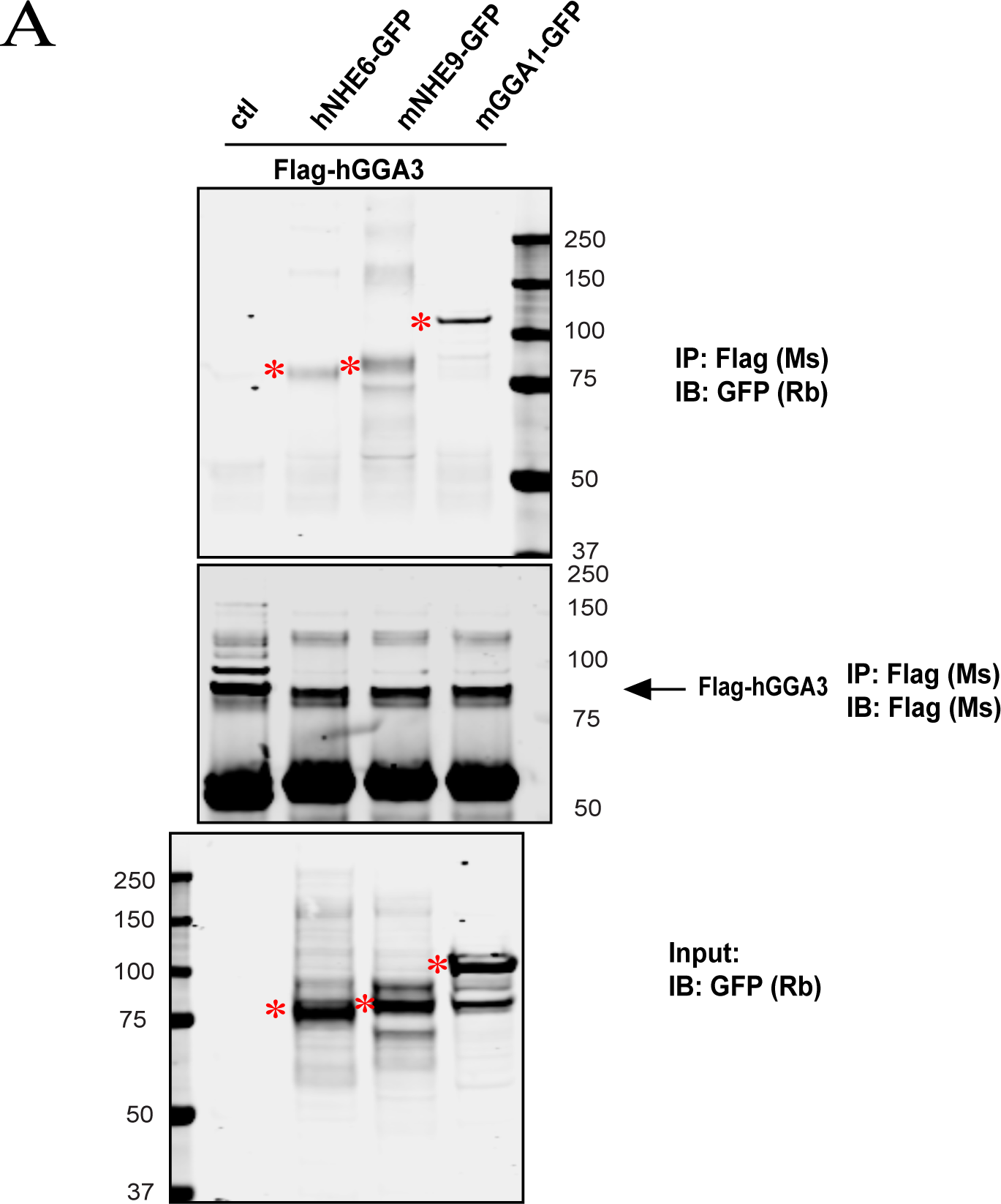
GGA3 interaction with NHE6 and NHE9. **(A)** Cell lysates from HEK293T cells expressing Flag-hGGA3 with GFP-tagged hNHE6, mNHE9, and mGGA1 were immunoprecipitated with anti-Flag antibody. The precipitates were probed with GFP and Flag antibodies. Western blot analysis was performed with anti-GFP to detect the expression of NHEs and mGGA1.

**Figure S3.**
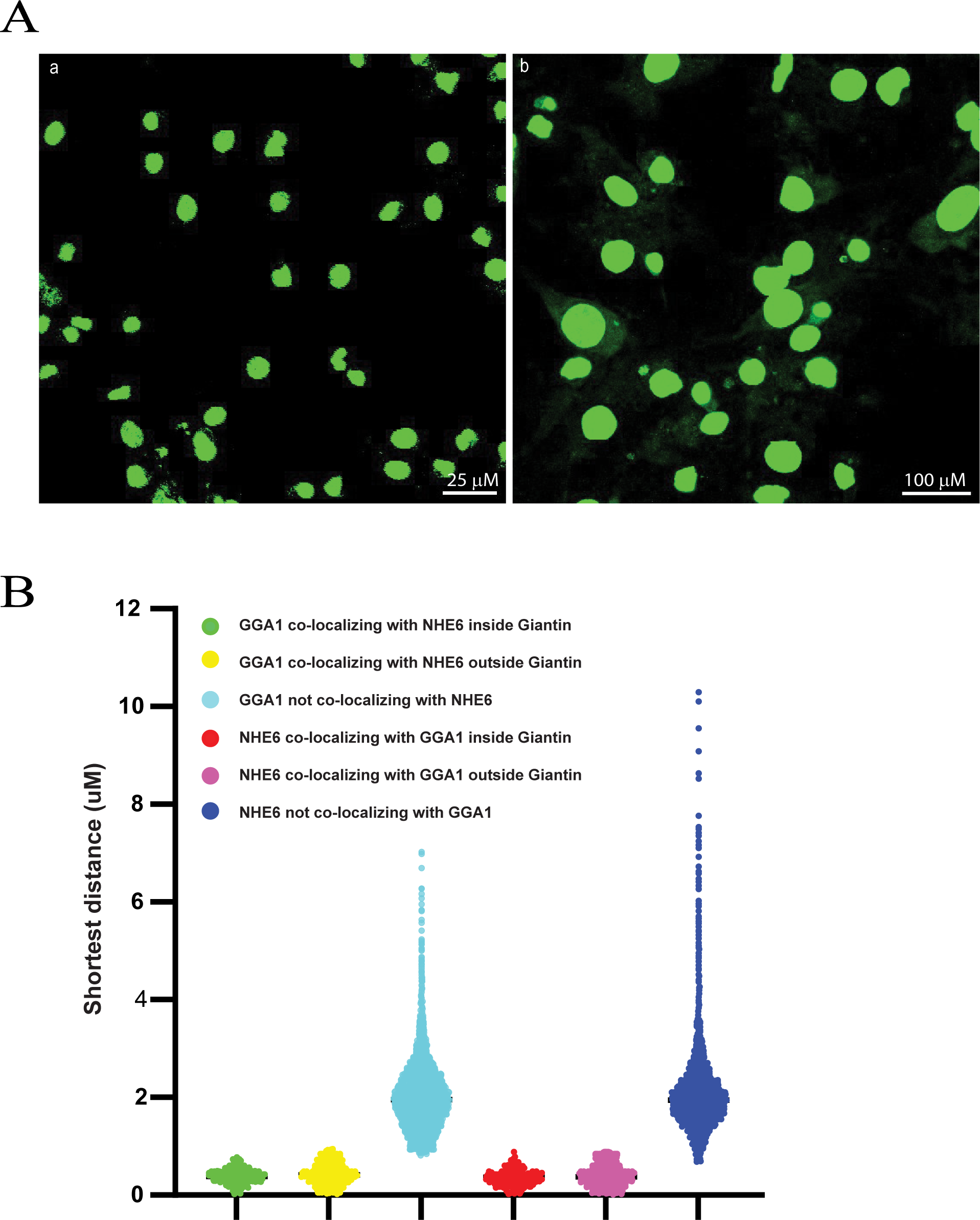
Shortest distance of NHE6 and GGA1 in rat primary hippocampal neurons. **(A)** Representative max projection z-stack images of unexpanded **(a)** or expanded **(b)** primary rat hippocampal neurons at 14 days *in vitro* (14 DIV) stained for nucleus using Sytox^TM^ Green Nucleic acid stain. **(B)** Measurement of the shortest distance of GGA1 to NHE6 or vice versa based on the classifications shown in Figure 5C. (n=14 neurons from 4 unique neuronal cultures).

**Figure S4.**
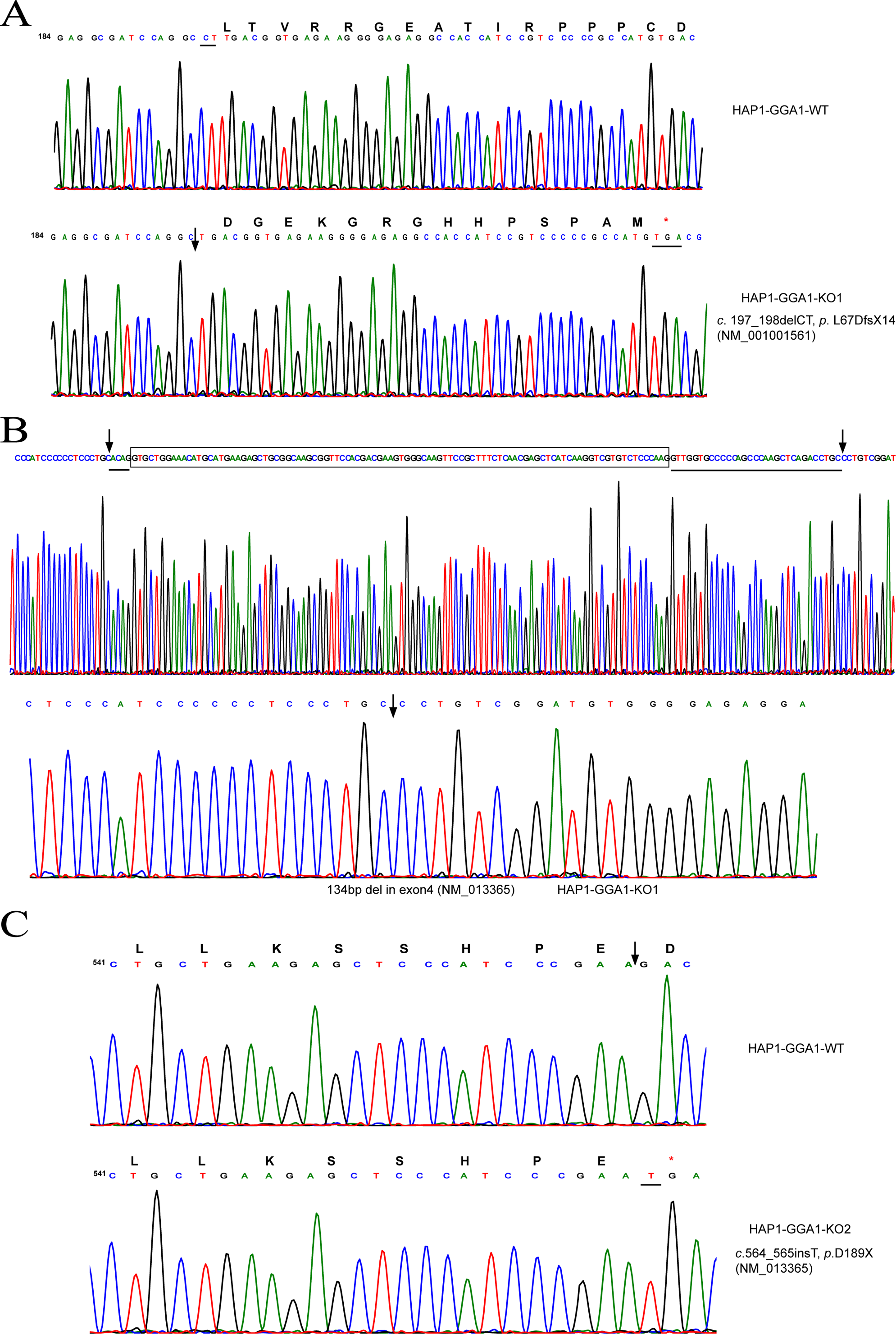
Sanger sequencing of HAP1 GGA1 Knockout (KO) cell lines. **(A, B)** Sanger sequencing from extracted DNA from HAP1 knockout (KO) and wildtype (WT) cell line PCR products. HAP1-GGA1-KO1, double knockout, has 2bp deletion in exon3 (*c*. 197_198delCT, *p*. L67DfsX14) (NM_001001561) (**A**) and 134bp deletion (**B**) in exon 4 (NM_013365). **(C)** HAP1-GGA1-KO2 has 1bp insertion in exon7 (*c*.564_565insT, *p*.D189X) (NM_013365).

**Figure S5.**
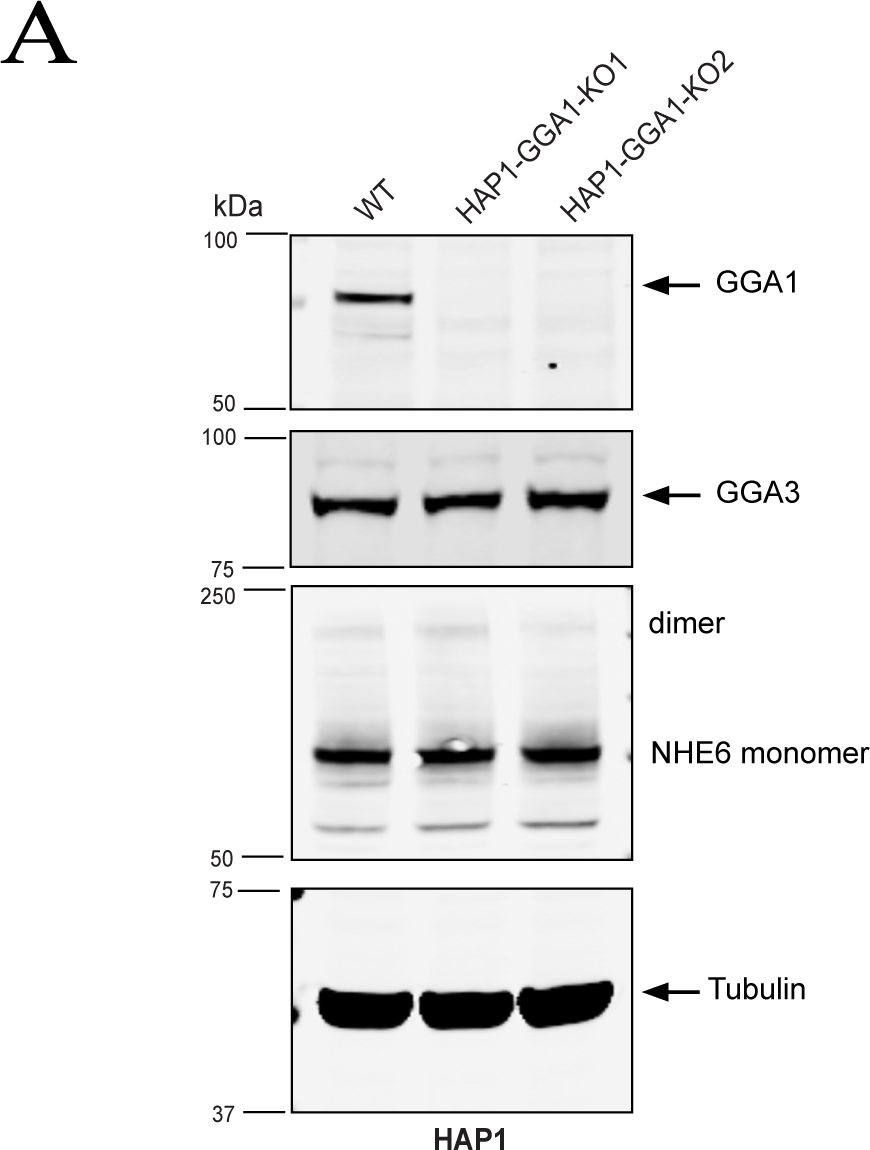
GGA1 Knockout (KO) lines do not express GGA1 protein. **(A)** Western blot analysis of cell lysates harvested from HAP1 knockout (KO) and wildtype (WT) cell lines using GGA1 antibody. Tubulin was used as a loading control and GGA3 was used for the specificity of GGA1 KO.

**Figure S6.**
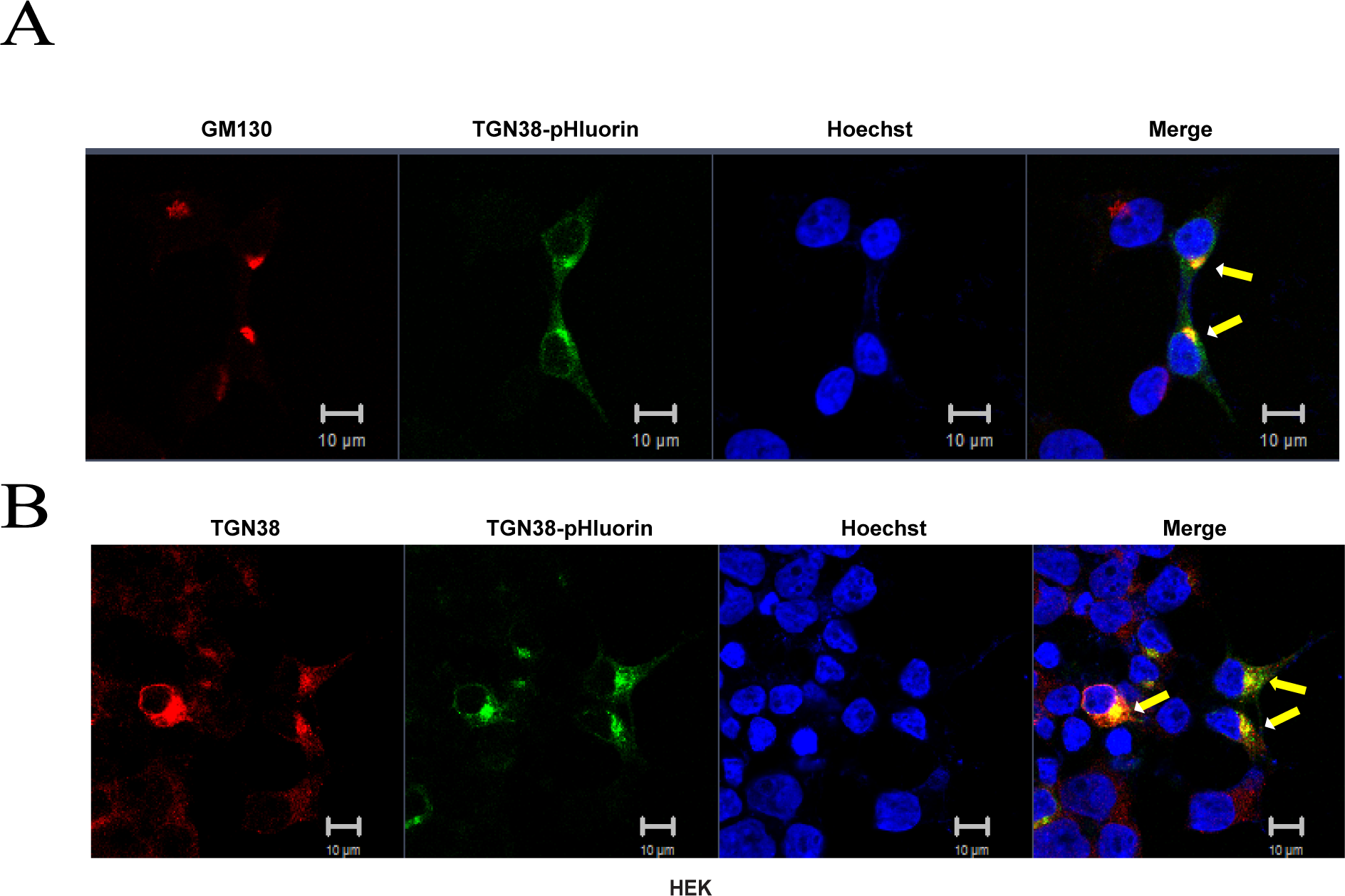
TGN38-pHluorin construct colocalizes with Golgi markers in HEK cells. Representative confocal microscopy images of HEK cells transfected with TGN38-pHluorin (green) and stained with (**A**) anti-GM130 (cis-Golgi) or (**B**) anti-TGN38 (trans-Golgi) antibodies (red). Hoechst=blue. Scale bar=10 μm.

---

**Table S1:**
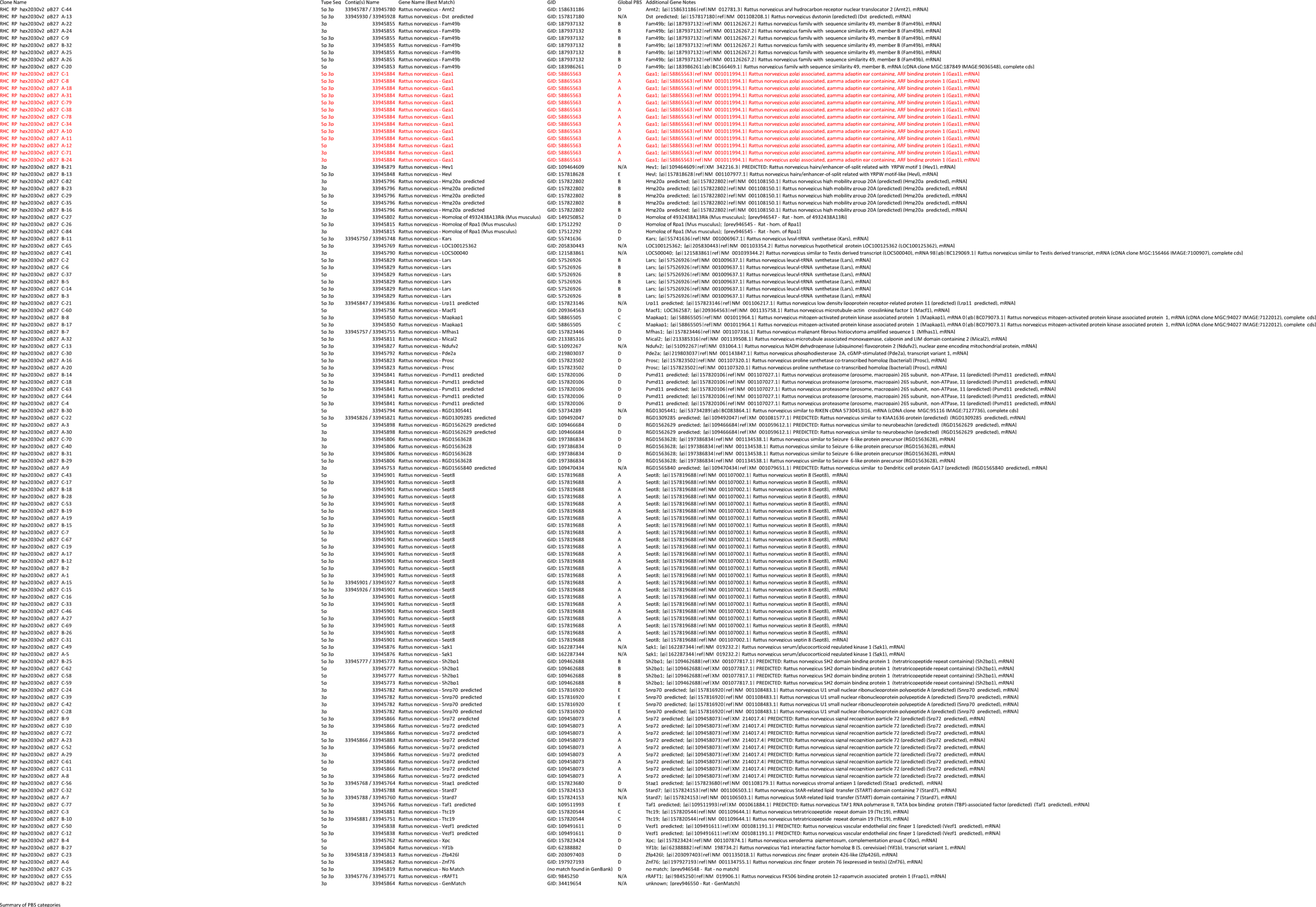
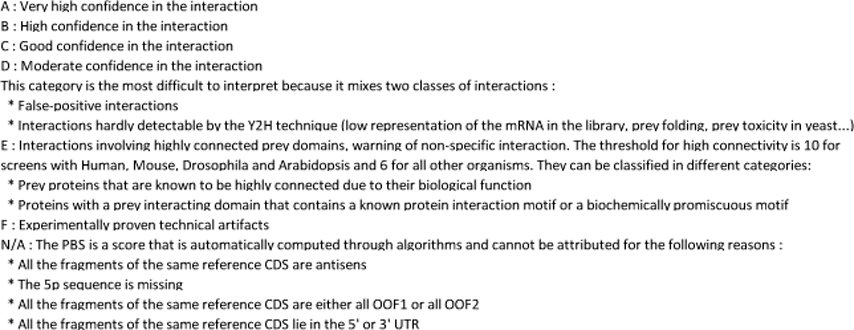
Mouse Slc9a6 (537-630) vs Rat Hippocampus RP1.

**Table S2:**
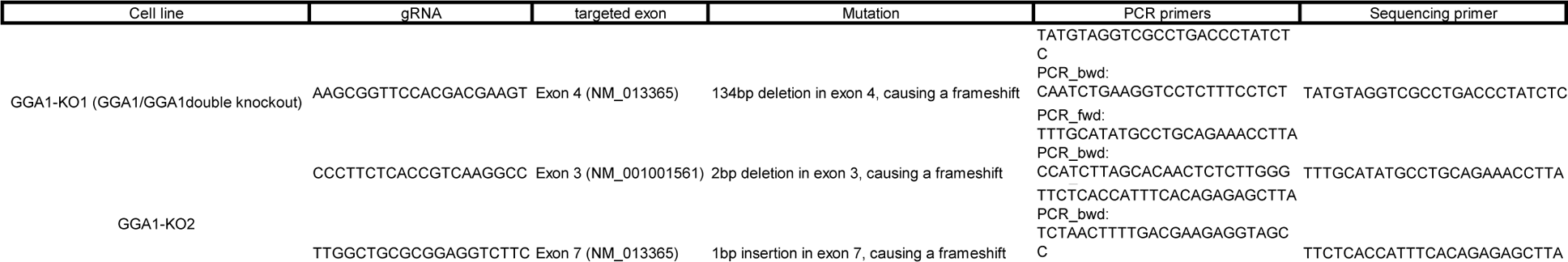
HAP1-GGA1 Knockout cell line Engineered using CRISPR/Cas9.

**Table S3:**
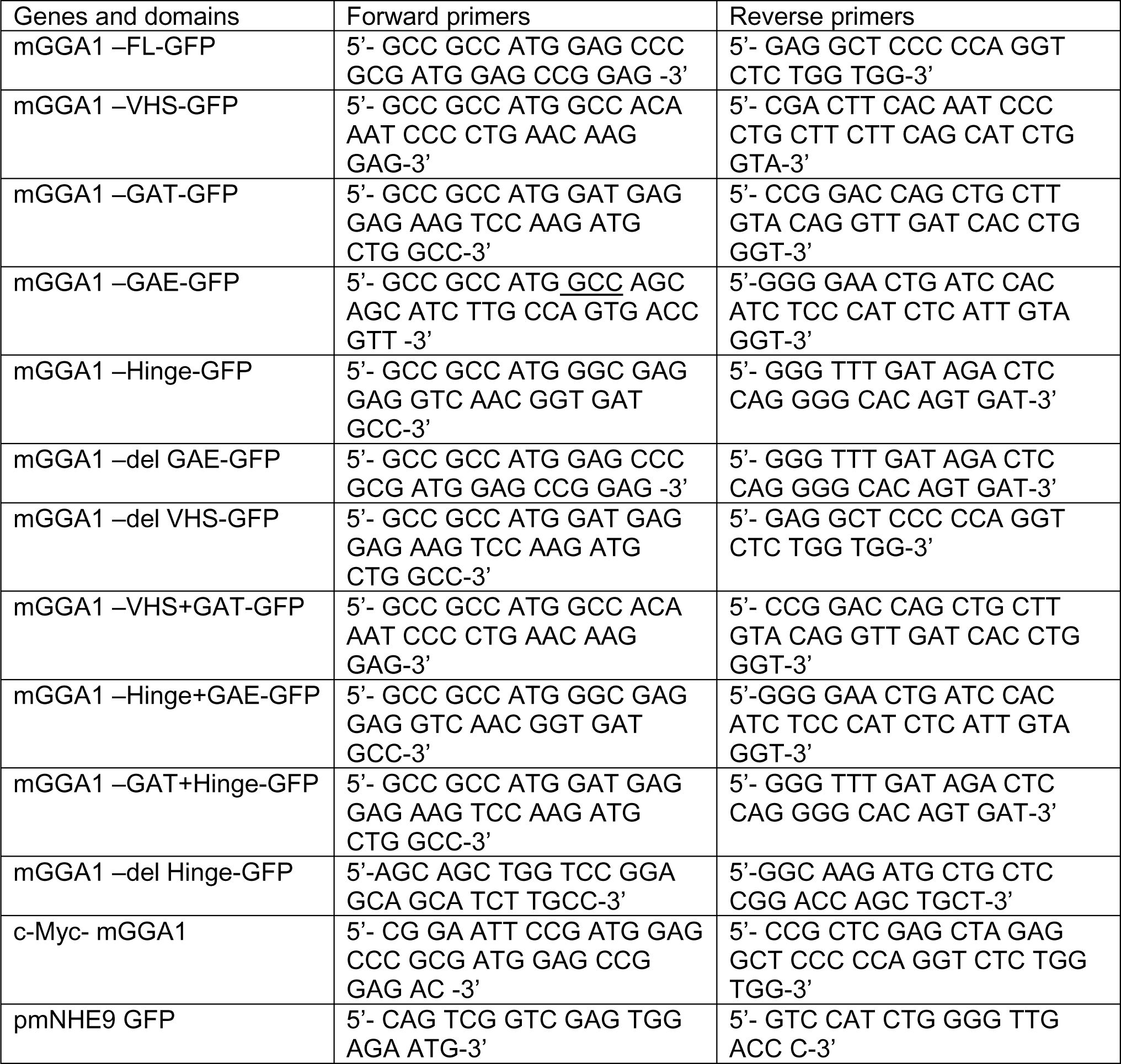
List of primers used for making GGA1, NHE9 constructs.

**Table S4:**
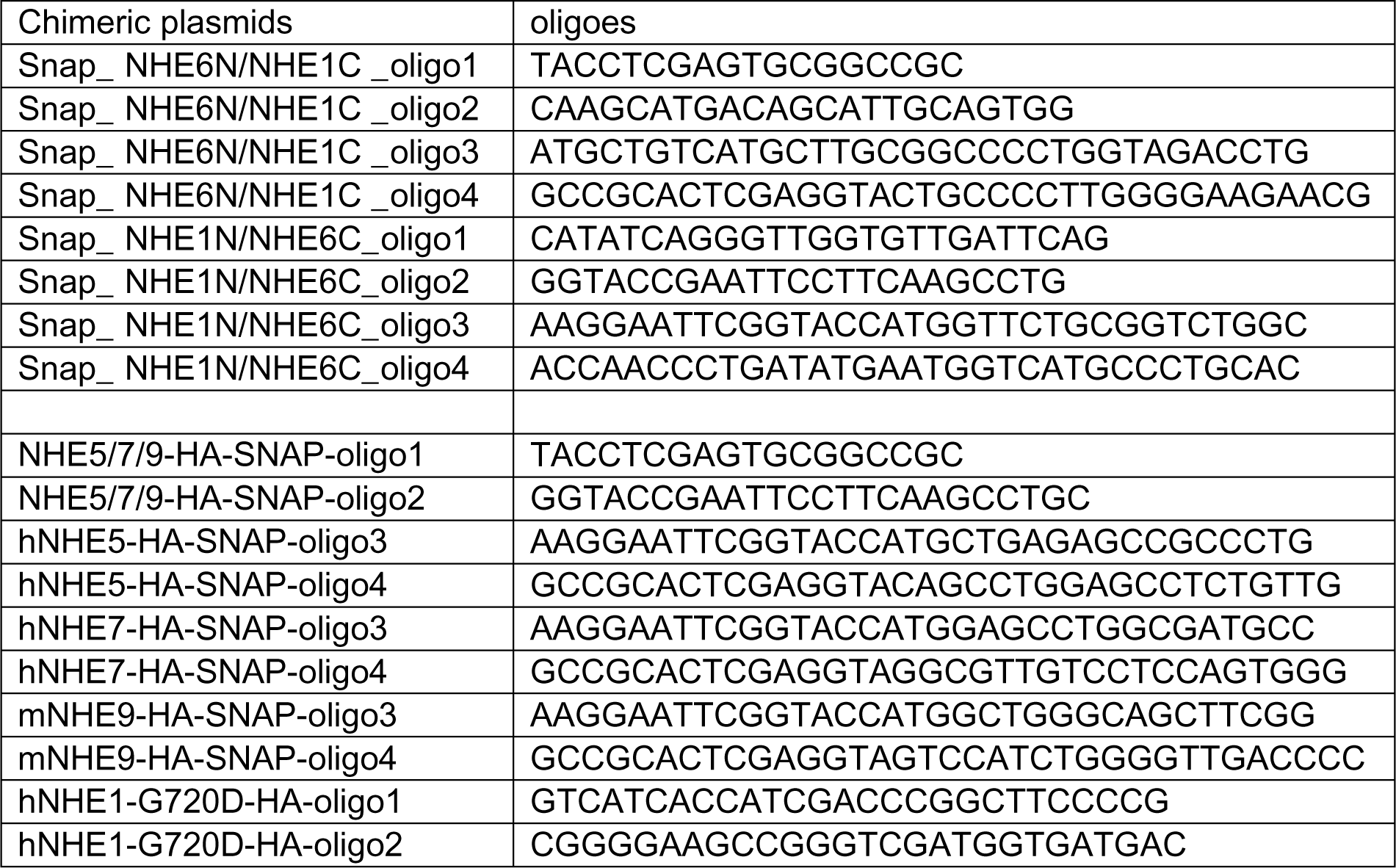
List of primers for making in fusion snap constructs.

**Table S5:** fractionation ultracentrifuge speed and time.

| Fractionation | Ultracentrifuge speed | Time (hrs) |
| --- | --- | --- |
| Endosome | 41,600 rpm | 3 |
| Lysosome | 145,000 g | 2 |
| Golgi | 120,000 g | 3 |

